# Structural and Computational Analysis of *Pseudomonas aeruginosa* DNA Gyrase Reveals Molecular Characteristics That May Contribute to Ciprofloxacin Resistance

**DOI:** 10.64898/2025.12.19.695517

**Authors:** Lalith Perera, Libertad Garcia-Villada, Andrea M. Kaminski, Natalya Degtyareva, Lars C. Pedersen, Paul W. Doetsch

## Abstract

*Pseudomonas aeruginosa* is considered a priority pathogen by the World Health Organization due to its resistance to antibiotics. Isolates resistant to ciprofloxacin (CPFX), a bactericide commonly used against *P. aeruginosa*, usually carry the mutations T83I or D87N in the GyrA subunit of the DNA gyrase. Yet, the molecular mechanisms by which these mutations confer CPFX-resistance to *P. aeruginosa* are unknown. Here we solved the crystal structure of the *P. aeruginosa* gyrase catalytic cleavage core and used it to carry out molecular dynamic (MD) simulations of CPFX-gyrase binding in the wild-type as well as the T83I and the D87N mutant systems. Our results show that DNA plays the most prominent role in maintaining the CPFX-bound conformation, with no appreciable contributions from Thr83 or Asp87. Interestingly, we found a solvent cavity adjacent to these residues that may provide CPFX access to the active site. Interaction energy analysis using Umbrella Sampling indicates that Thr83 and Asp87 may influence CPFX trajectory during binding. In the mutant systems, the attractive potential decreases, which may hinder CPFX accessing the binding site. These results shed light on *P. aeruginosa* resistance to CPFX and may help provide a methodology to identify new therapeutic agents to target fluoroquinolone resistant bacteria.

**Graphical abstract caption:** 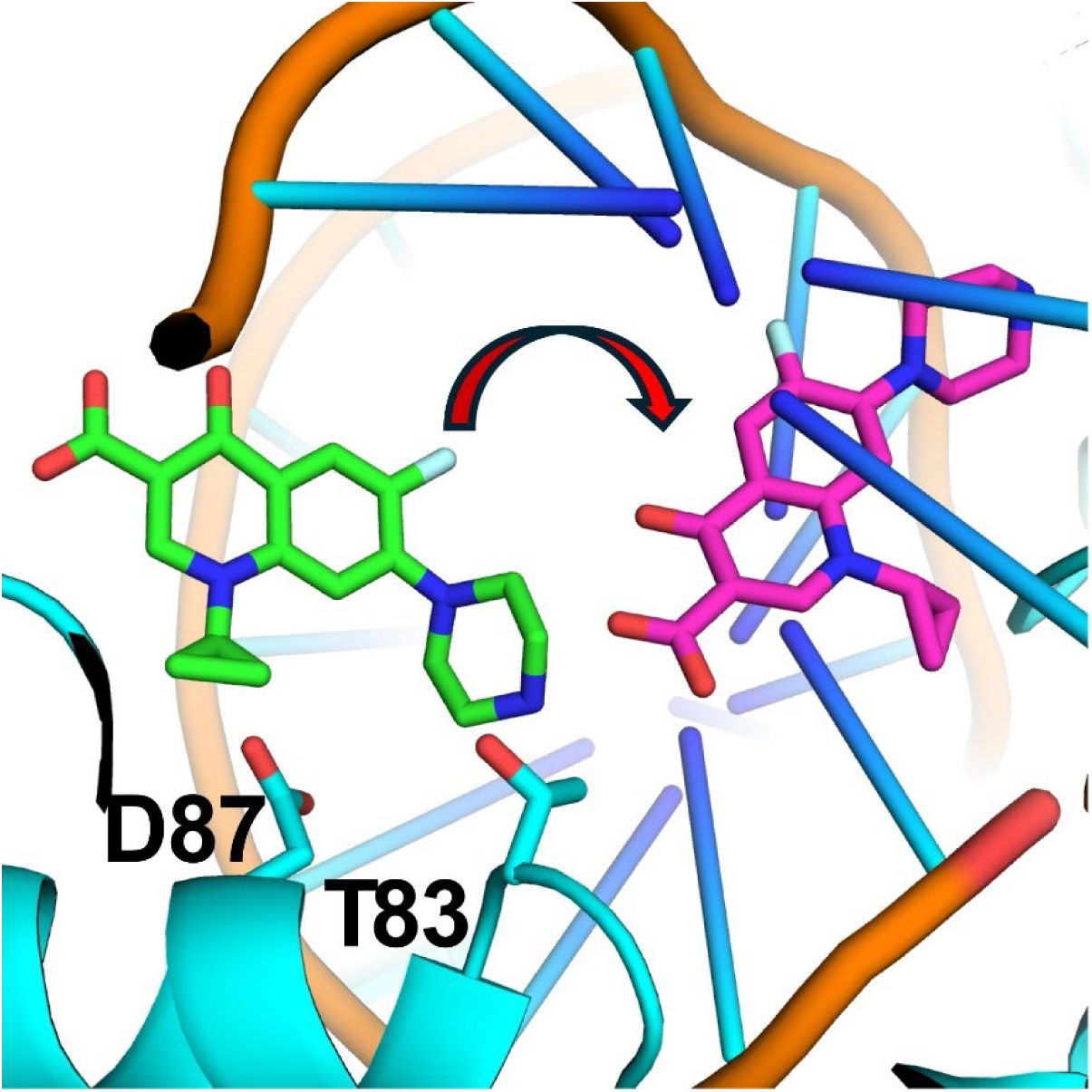

Molecular dynamic positioning of ciprofloxacin (CPFX) from the cavity site (green) to the bound inhibition site (magenta) suggests that mutations T83I and D87N, conferring CPFX resistance to the *P. aeruginosa* gyrase, may have an influence on the ability of CPFX to access the binding site. Protein is shown in cyan, DNA backbone in orange, and DNA bases in gradient from cyan to blue.

## 1. Introduction

*Pseudomonas aeruginosa*, a ubiquitous gram-negative bacterium, is an important opportunistic pathogen causing high morbidity and mortality. It is a major species contributing to nosocomial infections in patients with compromised immune systems, and the leading cause of chronic lung infections in cystic fibrosis and chronic obstructive pulmonary disease patients [1–3]. Infections are difficult to treat because *P. aeruginosa* is naturally resistant to high concentrations of many common, first-line antibiotics and has a remarkable ability to acquire mutations conferring resistance to antimicrobial agents [4–8]. The World Health Organization (WHO) has listed it as one of the six pathogens (***E****nterococcus faecium*, ***S****taphylococcus aureus*, ***K****lebsiella pneumoniae*, ***A****cinetobacter baumannii*, ***P****. aeruginosa*, and Enterobacter species or **ESKAPE** organisms) as top species prioritized for research due to their development of resistance to antibiotics [9].

Quinolones are one of the few antibiotic classes that are effective against *P. aeruginosa* infections, with the fluoroquinolone ciprofloxacin (CPFX) among the most used [10]. Their intense application, though, has led to a rapidly increasing bacterial resistance to this group of antibiotics and treatment failure [11–14]. Resistance is often determined by chromosomal mutations affecting drug permeability (increased efflux or reduced levels of drug accumulation in the cells) or type II topoisomerase (DNA gyrase, the quinolones’ target) sensitivity to the drug [11–14]. In CPFX-resistant isolates of *P. aeruginosa* identified in the clinic, the most frequently found mutations are mapped to the gene *gyrA*, which codes for the gyrase subunit GyrA (reviewed in [10]). Mutations in GyrA are located in a domain termed the “Quinolone Resistance-Determining Region” (QRDR), which encompass amino acids 67 to 106, with the most common being the specific mutations GyrA T83I and GyrA D87N [10,12,15].

DNA gyrase is an ATP-dependent bacterial type II DNA topoisomerase that performs the critical function of introducing negative supercoils into a double-stranded DNA (dsDNA) to prevent hyper-supercoiling. DNA gyrase is a heterotetrameric enzyme comprised of two heterodimers (complex I and II), each containing one GyrA and one GyrB subunit (Figure 1A). Both subunits are involved in the binding and cleaving of DNA; GyrB also hydrolyzes ATP. To produce negative supercoils, gyrase binds two close segments of dsDNA: a G (or Gate) segment, and a T (or Transported) segment (Figure 1A).

**Figure 1.**
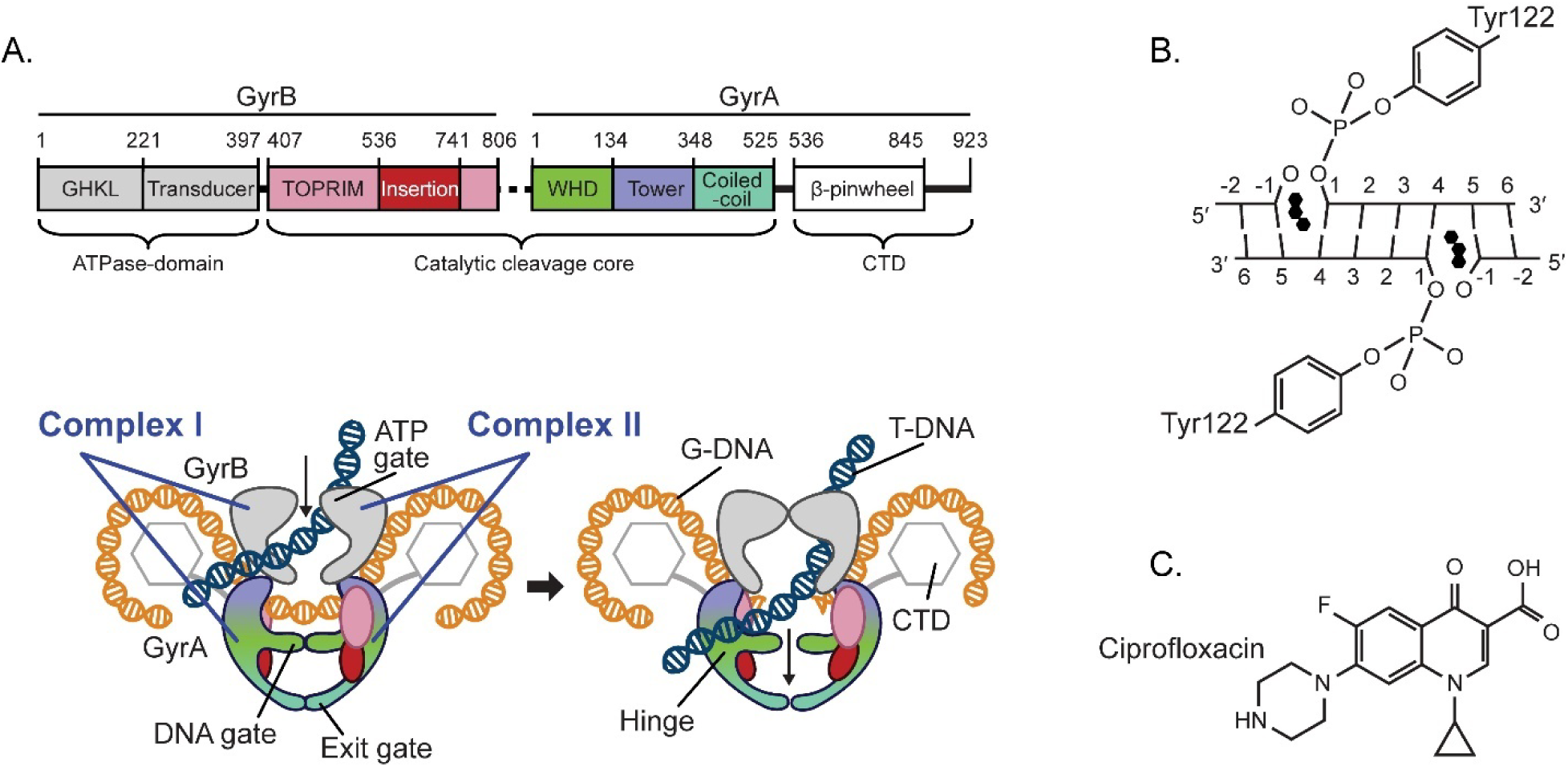
*P. aeruginosa* DNA gyrase structure and function. (A) Domain organization of the P. aeruginosa gyrase GyrB/GyrA complex with a cartoon depiction of the strand passage of the T-DNA segment through the temporarily cleaved G-DNA segment to introduce a -1 supercoil in the DNA. Colored regions represent the catalytic core expressed as in the GyrB-GyrA fusion protein used in this study. Dotted line in linear schematic represents the covalent linker (GGS) that connects GyrB to GyrA in our construct. CTD stands for C-Terminal Domain. (B) To initiate the DNA double strand break, the nucleophilic Tyr122 residue from each GyrA molecule in the complex attacks the phosphate backbone on opposite strands four base pairs apart, forming a covalent protein/DNA complex. Quinolones such as ciprofloxacin (CPFX, three black hexagons) can stack between the bases with high affinity at each break site, inhibiting passage of the T-DNA segment. (C) Molecular schematic of CPFX. (Figure inspired by Chan et al. [19]).

Then it cleaves both strands of the G-segment through nucleophilic attack by a tyrosine located in each of the GyrA subunits, leaving a four-base stagger. This results in the formation of a reversible covalent bond between the newly generated 5’-phophates and the catalytic tyrosine residues (Figure 1B). DNA cleavage is synchronized with ATP hydrolysis-facilitated passage of the T-segment through the G-segment and DNA-gate, followed by re-ligation of the G-segment via the reverse reaction and release of the DNA strands (reviewed in [16–18]).

Quinolones inhibit DNA gyrase through the formation of a stable ternary complex with the gyrase and covalently bound DNA whereby two quinolone molecules intercalate the DNA at both nick sites of the G-segment (Figure 1B), thus restraining the gyrase from re-ligating this segment (reviewed in [13,14,20–22]). These complexes can be reversed at low quinolone concentrations, but high drug concentrations produce dsDNA breaks that lead to chromosomal fragmentation and cell death (reviewed in [14,20]).

Analyses of wild-type and quinolone-resistant mutant gyrase cryo-EM and crystal structures alone or in combination with molecular docking and molecular dynamics (MD) simulations are useful means for understanding the drug-gyrase interaction and the drug mechanism of inhibition. Such structural studies provide the basis for the design of novel antibacterial agents or modification and implementation of existent ones (e.g. [19,21,23–28]). Partial or complete crystal or cryo-EM gyrase structures exist for *Escherichia coli* [29,30], *S. aureus* [19,31], *A. baumanii* [23], *Thermus thermophilus* [32], *Streptococcus pneumoniae* [33,34], and *Mycobacterium tuberculosis* [24], but not for *P. aeruginosa*.

To date, dynamic models of *P. aeruginosa* gyrase are based on the gyrase structures of other bacteria (e.g., [10,15,35]), which may render an incomplete picture of how resistant-mutant versions of this enzyme interact with different drugs.

Here, to understand the fluoroquinolone resistance mechanism in *P. aeruginosa* at an atomic level, we examine how the GyrA mutations T83I and D87N modify *P. aeruginosa* gyrase interactions with CPFX (Figure 1C). We generated a *P. aeruginosa* GyrB/GyrA fusion construct that concatenates the DNA binding/cleavage domains and solved its structure using crystallography in the apo- and DNA/CPFX-bound forms. These structures were used as starting models for MD analysis of wild-type (WT) and mutant proteins to examine how the T83I and D87N mutations may contribute to CPFX resistance. The combined structural/computational analysis revealed that these residues lie along a potential solvent entry channel providing CPFX access to the active binding site. How these residues impact access and may affect resistance is discussed.

## 2. Materials and Methods

### 2.1. Cloning, Expression, and Purification of P. aeruginosa Gyrase Catalytic Cleavage Core

A construct (referred to as *gyrB*-GGS-*gyrA525*) composed of full-length *P. aeruginosa gyrB* (coding for residues 1 to 806)-linker (GGS)-truncated *gyrA* (coding for residues 2 to 525), optimized for expression in *E. coli*, was obtained through gene synthesis (Genewiz). A truncated *gyrB* (coding for residues 397 to 806)-linker (GGS)-truncated *gyrA* (coding for residues 2 to 525) construct (referred to as *gyrB397*-GGS-*gyrA525*) (Figure 1A) was amplified using the synthetized version as template and the following primers: forward primer, 5′-TCGgcggccgcAATGAAGGGTGCGCT-3′; reverse primer, 5′-TAGaagcttTTAGCTGGCCACAAT-3′. (Lowercase letters denote NotI and HindIII restriction sites, respectively). Both constructs were cloned into a pGEXM vector using the NotI/HindIII restriction sites, and sequenced. pGEXM vectors carrying the constructs were transformed into Rosetta2(DE3) cells for expression. Overnight (ON) cultures were grown at 37°C in shaker flasks in Terrific Broth (TB) media containing 100 µg/ml ampicillin and 35 µg/ml chloramphenicol. The next day, the ON cultures were added to Fernbach flasks containing 1 L of TB media and antibiotics and placed at 37°C on shaker.

When the OD600 reached 0.9, the temperature was set to 18°C. After 45 minutes, IPTG (isopropal-β-D-1-thiogalactopyranoside) was added to a concentration of 200 µM and the cultures were shaken ON at 18°C. Cells were then pelleted by centrifugation at 4000 g for 15 min and resuspended in sonication buffer consisting of 25 mM Tris pH 7.5, 500 mM NaCl, 1 mM EDTA, and 1 mM DTT, followed by sonication. Cell debris were pelleted by centrifugation for 35 minutes at 47,900 g. The soluble fraction was equilibrated with Glutathione 4B Sepharose resin (Cytiva) in batch on a rocker at 4°C for 1.25 hours and the resin was washed multiple times in batch with sonication buffer. Protein was removed from the resin bound glutathione-S-transferase with TEV (Tobacco Etch Virus) in batch on rocker at 4°C ON. Protein was concentrated, then further purified by gel filtration on a Superdex200 gel filtration column (Cytiva) equilibrated in 25 mM Tris pH 7.5, 75 mM NaCl, 1 mM DTT. Eluted fractions containing protein were pooled and concentrated. While the *gyrB*-GGS-*gyrA525* construct produced little protein, the *gyrB397*-GGS-*gyrA525* overexpressed well and could be concentrated to ∼19 mg/ml for crystallization experiments.

### 2.2. Crystallization and Structure Determination of P. aeruginosa Gyrase Catalytic Cleavage Core

Crystals of the apo *gyrB397*-GGS-*gyrA525* protein were grown at 4°C using the vapor diffusion hanging drop method. This was originally an attempt to obtain a protein/DNA complex as the protein used consisted of 17.9 mg/ml protein mixed with an annealed 24mer duplex DNA (2:1 DNA to protein ratio; 24merF GGTCATGAATGACTATGCACGTAA, 24merR TTACGTGCATAGTCATTCATGACC) in 25mM Tris pH 7.5, 150 mM NaCl, and 5 mM MgCl2, that was allowed to equilibrate for 4 hours at 4°C prior to setup. For crystallization, a solution consisting of 2 µl of protein/DNA complex was mixed with 2 µl of the reservoir consisting of 100 mM HEPES pH 7.5, 5% PEG 8000 (Microlytic), and 26% MPD (Hampton Research) equilibrated for one day against the reservoir prior to streak seeding. The crystal used for data collection was harvested after six days, transferred to a cryo-solution consisting of 100 mM HEPES pH 7.5, 40% MPD, and 5% PEG 8000, then flash frozen in liquid nitrogen. Data were collected on the Berkeley Center for Structural Biology beamline 5.0.2 at The Advanced Light Source, and processed using HKL3000 [36]. Protein Data Bank coordinates (PBD) 3NUH were used as a search model for molecular replacement in PHASER [37,38].

The model was refined with multiple cycles of refinement in PHENIX and manual model building in COOT [39–41]. No DNA was visible in the electron density. Several regions within the structure lack sufficient electron density to be modeled: GyrB residues 397-403, 453-461, 503-504, 570-575, 703-706, 717-720, and 798-806; GGS linker; GyrA residues 2-5, 175-176, 401-405, 415-420, and 427-430). Data statistics and model analysis are shown in Table I.

**Table I.**
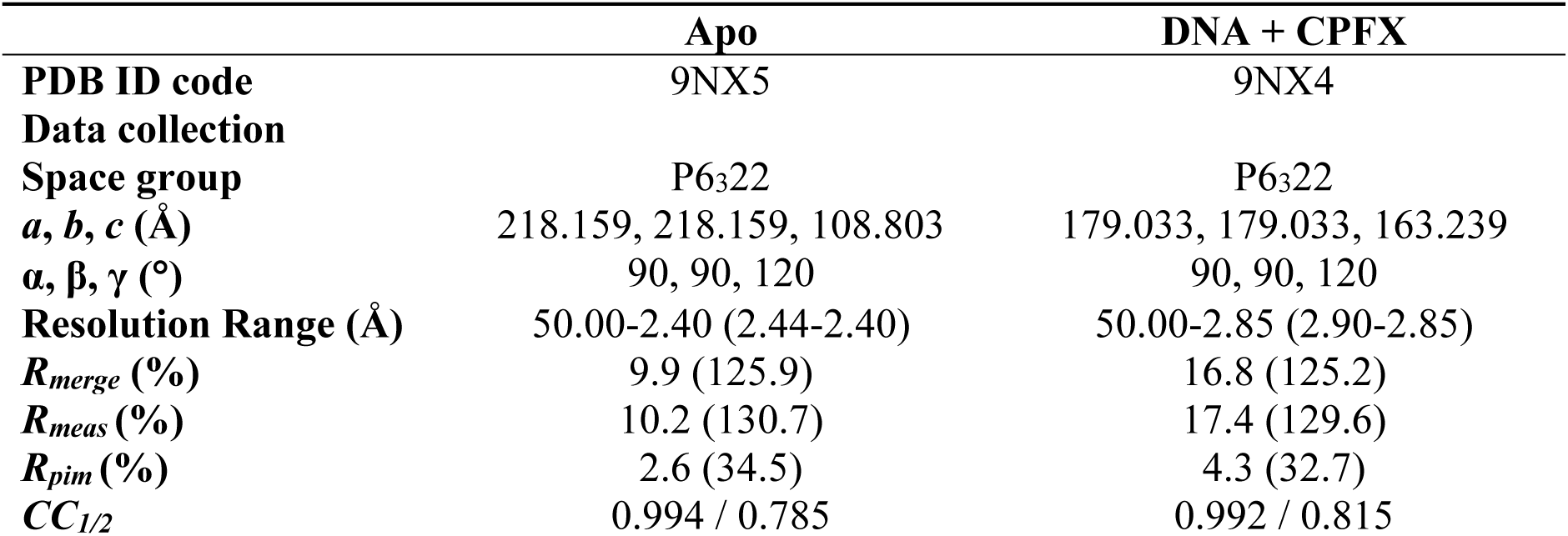
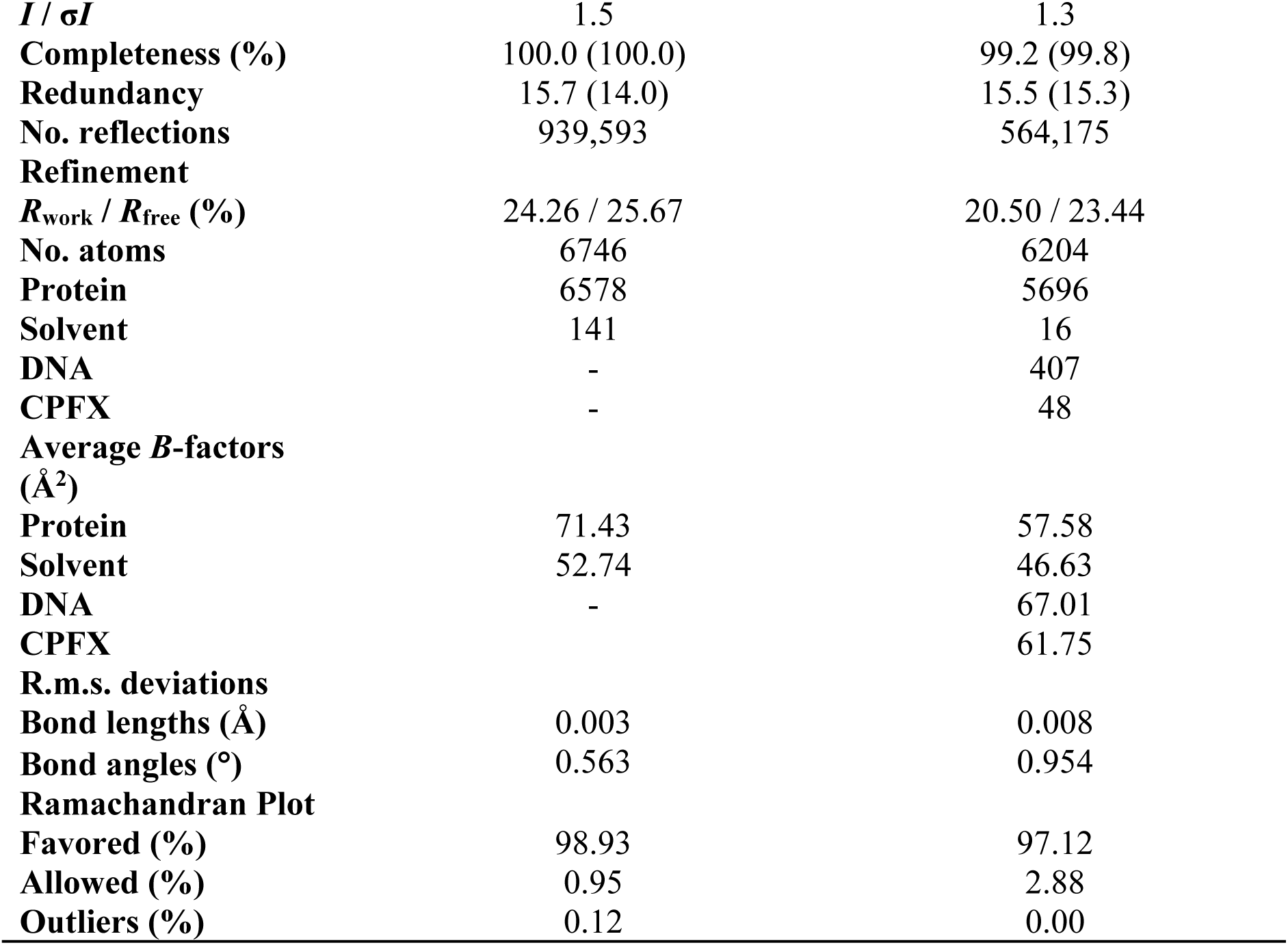
Crystallographic data collection and refinement.

Crystals of the DNA\CPFX bound *gyrB397*-GGS-*gyrA525* protein were grown at 21°C using the vapor diffusion hanging drop method. To form the complex, 16 mg/ml of protein was equilibrated with an annealed 20mer duplex DNA (3:1 duplex DNA to protein ratio; 20mer AGCCGTAGGGCCCTACGGCT) in 25mM Tris pH 7.5, 75 mM NaCl, 1 mM MnCl2, 1 mM DTT, and 1 mM CPFX, and allowed to equilibrate for ∼5 hours. For crystallization, 2 µl of protein/DNA/CPFX complex were mixed with 1 µl of reservoir solution consisting of 100 mM sodium acetate pH 6.0, 200 mM ammonium sulfate, and 14% PEG 4000 (Microlytic), and equilibrated for one day against the reservoir prior to streak seeding. Crystal was harvested after 11 days and flash frozen in liquid nitrogen in a cryo-solution consisting of 100 mM sodium acetate pH 6.0, 200 mM ammonium sulfate, 15% PEG 4000, 20% ethylene glycol, and 1 mM CPFX. Data were collected on the APS SER-CAT 22ID beamline at Argonne National Laboratory. The apo model of *gyrB397*-GGS-*gyrA525* was used to solve the molecular replaced for the DNA\CPFX bound structure using PHASER [37]. The DNA and CPFX were built into the map, and the model was refined with multiple cycles of refinement in PHENIX and manual model building in COOT [39–41]. The following residues lack sufficient density to be modeled: GyrB 397-404, 479-482, 562-735 (insertion domain), and 799-806; GGS-linker; GyrA 2-6. Quality of both the apo and DNA bound structures were assessed using MolProbity [42]. The apo and DNA bound structures have been deposited in the PDB with ID codes 9NX5 and 9NX4, respectively, with data statistics and model analysis results reported in Table I.

### 2.3. Determination of P. aeruginosa Gyrase Catalytic Cleavage Core Construct Activity

Determination of *gyrB397*-GGS-*gyrA525* activity and inhibition by CPFX was performed according to Nitiss et al., 2012 [43]. In brief, 20 µl reaction mixtures were prepared with 11 µl of H2O, 2 µl of 10 x buffer (ProFoldin *P. aeruginosa* DNA gyrase assay kit plus-100; final concentrations are: 20 mM Tris-HCl pH 8, 35 mM NH4OAc, 4.6% glycerol, 1 mM DTT, 0.005% Brij35, 8 mM MgCl2), 1 and 2 µl of purified *gyrB397*-GGS-*gyrA525* (1 and 2 µM final concentrations), 1µl of pUC18 (30 nM final concentration), 2 µl of 10 mM ATP (1 mM final concentration), and 2 µl of either H2O or CPFX solution (final concentrations 2, 4, and 8 µg/ml). Positive controls included two reactions with commercial *E. coli* gyrase (Sigma; 1 and 2 µl of a 2 units/µl solution), and negative controls included all components except the gyrase. Reaction mixtures were incubated at 37°C for 1 hour and stopped by adding 5 µl of 5 x stop solution (Topogen; 5% Sarkosyl, 0.125% bromophenol blue, 25% glycerol).

Proteinase K solution (Invitrogen; 20 mg/ml) was then added to a final proteinase concentration of 0.4 mg/ml, and the reaction mixtures were incubated at 37°C for 30 minutes. A sample (20 µl) of the mixtures was loaded in a 0.8% agarose gel. After electrophoresis, the gel was stained for 30 minutes in an ethidium bromide solution before visualization (Figure S1).

### 2.4. Molecular Dynamics (MD) Simulations

The initial structures of GyrB (residues 405–561 and 736–798) and GyrA (residues 7–525) of *P. aeruginosa* were derived from the X-ray crystal structure (PDB ID 9NX4). Missing residues were added using Modeller-10.2 [44]. DNA nucleotides spanning the nick sites (Figure 1B) were introduced based on the structure of *M. tuberculosis* (PDB ID 5BTC) [24]. CPFX molecules were realigned, and magnesium ions were incorporated as in the crystal structure mentioned above.

To prepare the starting configurations for MD trajectories, missing atoms and protons were added using the LEaP module of Amber.22 [45]. Counter ions were introduced, and the system was solvated in a box of TIP3P water, with the box boundary extending 20 Å from the nearest peptide atom. This setup resulted in a simulation box containing 279,472 atoms. Histidine residues 132, 195, 358, 379, and 465 in GyrB were δ-protonated, while the remaining histidine residues were ε-protonated. Charge neutralization and a 100 mM effective salt concentration, approximating physiological conditions, were achieved by adding 359 Na⁺ and 231 Cl⁻ ions.

Before equilibration, the solvated system underwent the following sequential steps:

1. **Belly dynamics**: 500 ps of simulation with fixed peptide.
2. **Energy minimization**: 5,000 steps.
3. **Low-temperature dynamics**: 1 ns of constant-pressure simulation at 200 K with a fixed protein, ensuring a reasonable starting density.
4. **Second energy minimization**: 5,000 steps.
5. **Stepwise heating**: Constant-volume MD simulations increasing the temperature from 0 K to 300 K over 3 ns.
6. **Constrained equilibration dynamics**: 20 ns of constant-volume simulation with a 10 kcal/mol constraint force applied only to backbone heavy atoms.

After a 30 ns equilibration period during which all constraints were removed, sampling was extended through three independent, constant-temperature, constant-volume MD simulations of 1 μs each. All MD trajectories were computed using the PMEMD (Particle Mesh Ewald Molecular Dynamics) module of Amber.22 with a 1 fs time step. Long-range Coulombic interactions were calculated using the PME method with a 10 Å cutoff for direct interactions. Amino acid parameters were derived from the FF14SB force field of Amber.22. CPFX atomic charges were calculated using ChelpG procedure of Gaussian.16 [46].

Configurations selected at each nanosecond of the production runs were used for all analyses utilizing the Cpptraj module of Amber.22 [47]. Interaction-free energies between gyrase residues and CPFX were calculated using the MM/PBSA (Molecular Mechanics/Poisson-Boltzmann Surface Area) protocol of Amber.22 [48]. These calculations were based on 1,000 configurations extracted at each nanosecond interval from each trajectory, resulting in a total of 3,000 configurations. An ionic strength of 0.1 M was selected for the MM/GBSA (Molecular Mechanics/Generalized Born Surface Area) calculations. Videos of the trajectories were generated using Chimera-1.18 [49].

Sets of MD runs were conducted for the WT as well as for the T83I and D87N mutant systems, with and without CPFX. The original WT simulation system was modified to construct these mutant systems. MD simulations were performed in triplicates, and the analysis methodology mirrored the approach described above for the wild-type system. Table S1 summarizes information on all systems studied in this work.

MD simulations in combination with Umbrella Sampling were carried out to evaluate the potential effects of specified mutations on CPFX binding. An area, identified from the original simulations, involving residues D87 and T83, referred to now as “the cavity site”, was deemed crucial as the entry point for CPFX along its transition pathway. CPFX was driven out from its binding site towards the cavity site by selecting 35 windows with 0.4 Å width along the direction from the bound position of CPFX towards the cavity site. For each window, a 1.2 ns of MD run was performed with the anchor strength of 100 kcal/mol. To apply the forces, a line along the center of mass of CPFX and the center of mass of four phosphates from the DNA that reside next to the cavity site was selected. Five hundred configurations selected from the last 1 ns of each window were used in the MM/GBSA energy analysis.

## 3. Results

### 3.1. Structural Analysis of P. aeruginosa Gyrase Catalytic Cleavage Core

Previous structural studies on bacterial DNA gyrases have demonstrated the benefits of fusing the N-terminus of GyrA to the C-terminus of GyrB [22,24]. In this report, we demonstrate that a *P. aeruginosa* gyrase catalytic cleavage core construct composed of residues 397-806 of GyrB fused with residues 2-525 of GyrA via a GlyGlySer linker (*gyrB397*-GGS-*gyrA525)* (Figure 1A) is functional, i.e. can generate DNA double strand breaks (Figure S1). To better understand how the T83I and D87N GyrA mutations may contribute to CPFX resistance in *P. aeruginosa*, we solved the structure of the *gyrB397*-GGS-*gyrA525* construct with and without DNA/CPFX bound (Figure 2A and C). The apo form crystallized with one molecule in the asymmetric unit forming the expected dimer with a neighboring symmetry-related molecule in the crystal (Figure 2A). While these structures are the first reported for *P. aeruginosa*, ample gyrase structural data exists for *E. coli*, which shares with *P. aeruginosa* ∼72% sequence identity within the gyrase catalytic cleavage core included in our construct [30,38,50]. Based on superposition, the apo catalytic cleavage core from both enzymes displays good agreement in the positions of the individual domains except for the domain unique to gram-negative bacteria, the insertion domain, that in *P. aeruginosa* pivots by ∼23° from the connection to the TOPRIM domain, resulting in a shift of greater than 20 Å for the remote residues (Figure 2B) [38].

**Figure 2.**
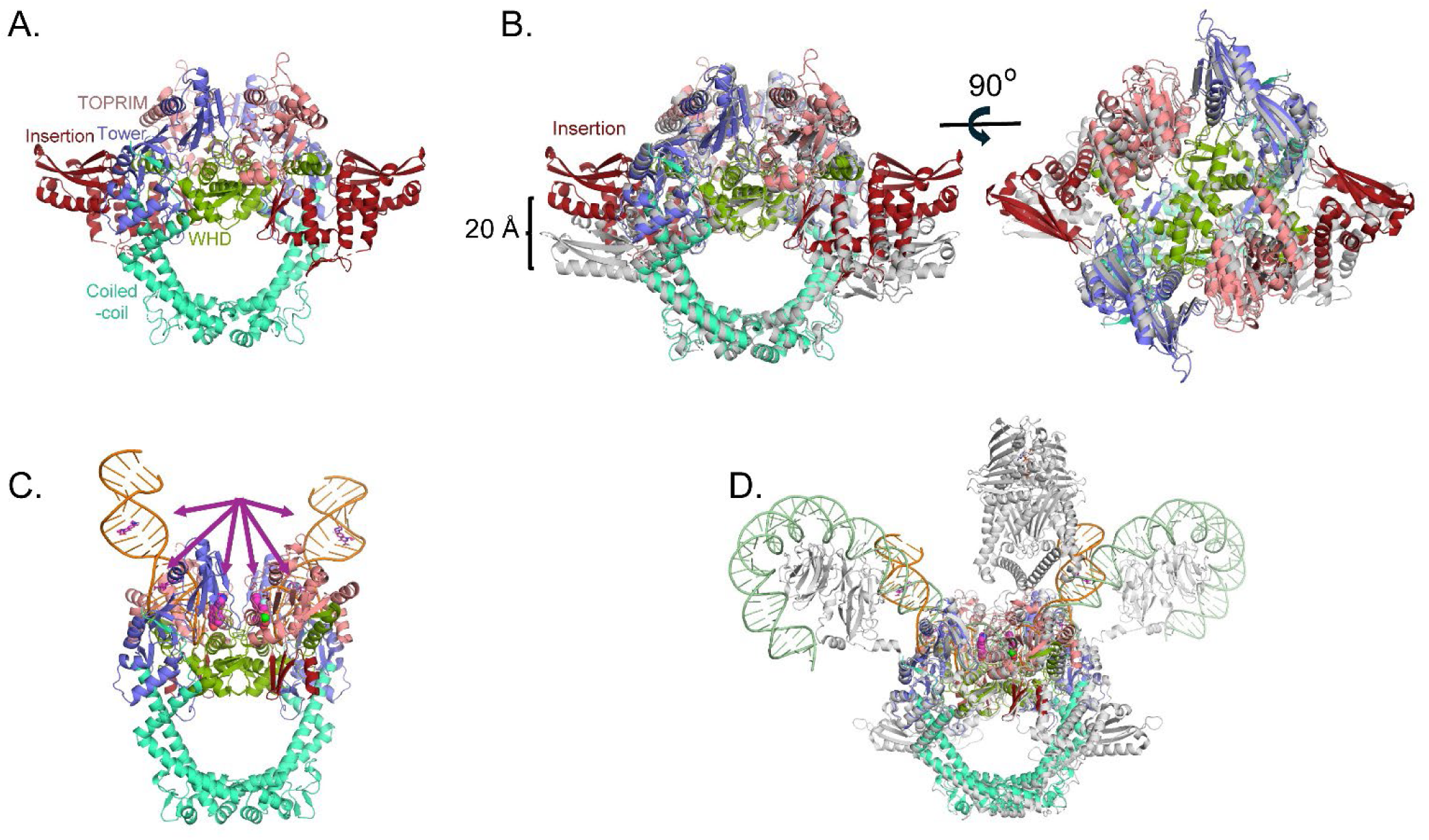
Crystal structure of *P. aeruginosa* gyrase catalytic cleavage core complex. (**A**) Side view of the apo dimer structure of the catalytic cleavage core. Domains are colored as in Figure 1A. (**B**) Two views of the superposition of the *P. aeruginosa* catalytic cleavage core with the apo *E. coli* catalytic cleavage core (gray; PDB ID 3NUH) [38]. The 20 Å separation of the insertion domain between the structures due to a 23° rotation is noted. (**C**) Structure of the *P. aeruginosa* DNA (orange) bound dimer catalytic cleavage core in complex with CPFX (magenta space filling). The arrows highlight CPFX binding positions in the crystal structure dimer. Based on crystal packing, a symmetry molecule of the catalytic core binding a ssDNA and CPFX, as well as ssDNA from two other asymmetric units, were used to generate the complex in this figure. In the structure, CPFX molecules are also found intercalating the duplex DNA outside the binding pocket (shown in stick; active site CPFX are shown in spheres). (**D**) Superposition of the full-length DNA bound form of the *E. coli* gyrase as determined by CryoEM (protein gray, DNA light green; PDB ID 6RKW [30]) onto the DNA bound form of *P. aeruginosa*. Superpositions between *E. coli* and *P. aeruginosa* illustrates similar conformations in the apo (**B**) and DNA binding structures (**D**) between the two enzymes.

Attempts were made to crystalize *gyrB397*-GGS-*gyrA525* with a duplex DNA substrate spanning the active site of the dimer utilizing 20mer and 24mer duplexes as had been done previously for other gyrase/topoisomerase II crystal structures [22,24]. Crystals were obtained in the presence of CPFX and a 20mer palindromic DNA sequence that resulted in one strand of DNA and one molecule of the catalytic cleavage core (GyrB/GyrA) in the asymmetric unit, forming the expected dimer with a crystallographic symmetry mate (Figure 2C). In this crystal structure, most of the insertion domain is disordered.

Interestingly, rather than obtaining the expected structure of the duplex DNA oligo binding across the active site of the two molecules in the crystallographic dimer, each active site is binding to the blunt end of the duplex DNA (Figure 2C and 3A). This DNA bound form of *P. aeruginosa’s* catalytic cleavage core also shows good agreement with the CryoEM structure of the full length *E. coli* gyrase that includes the N-terminal ATPase domain of GyrB and the C-terminal pinwheel domain of GyrA in a closed conformation binding DNA (Figure 2D) [30]. While the blunt end superimposes well with the position of DNA in other gyrase structures such as in *E. coli*, *S. aureus* and *M. tuberculosis* (*Mtb*), it leaves a gap that could accommodate four nucleotide base pairs between the two ends, consistent with spacing in other structures (Figure 4A) [22,24,30]. For *P. aeruginosa*, CPFX is found stacking with the blunt end of the DNA on one side and a solvent citrate molecule positioned on the other (Figure 4A and B). While the location of CPFX binding to the complex is similar to what has been seen in the *S. aureus* and *Mtb* structures, the relative orientation of CPFX is different, possibly due to the lack of duplex DNA on both sides of the drug (Figure 4A) [22,24].

**Figure 3.**
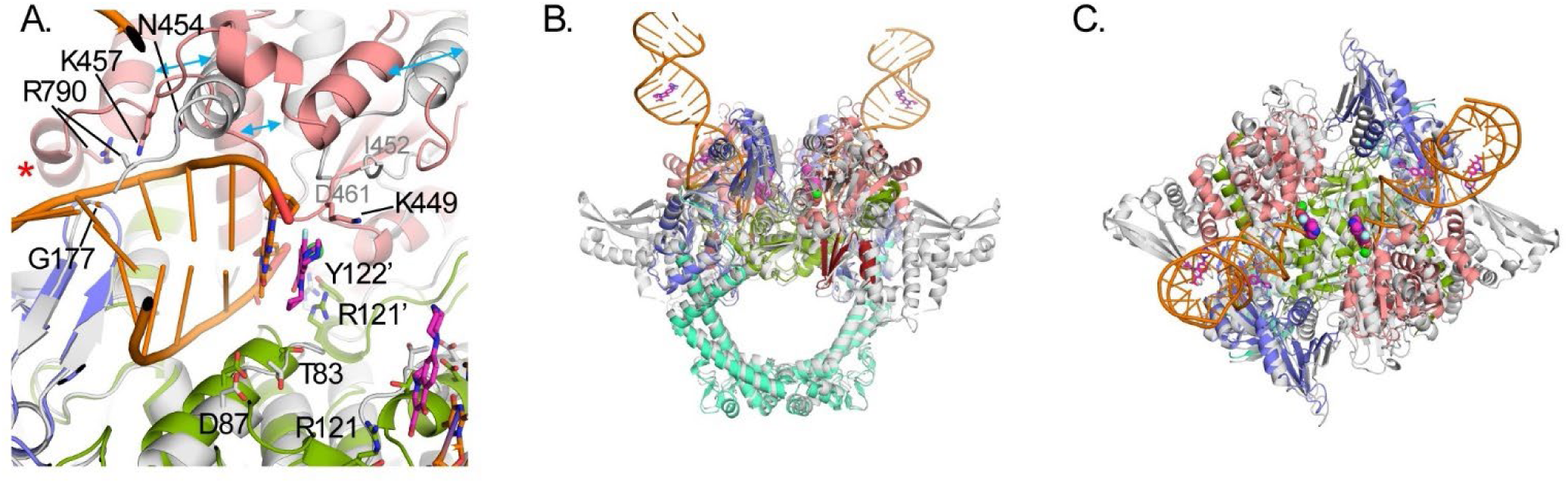
Comparisons of the apo and DNA-bound forms of *P. aeruginosa* gyrase catalytic cleavage core. (**A**) Superposition of the DNA binding site from the apo (colored gray) and DNA-bound form (colored as in Figure 2A) of the *P. aeruginosa* gyrase. The DNA binds with a blunt end in each active site. In the DNA bound form, the C-terminal region of GyrB containing residue Arg790 (indicated by red asterisk) is displaced from the DNA binding site. (**B**) and (**C**) Side and top views of the superposition between apo and DNA/CPFX bound structures of the *P. aeruginosa* gyrase.

**Figure 4.**
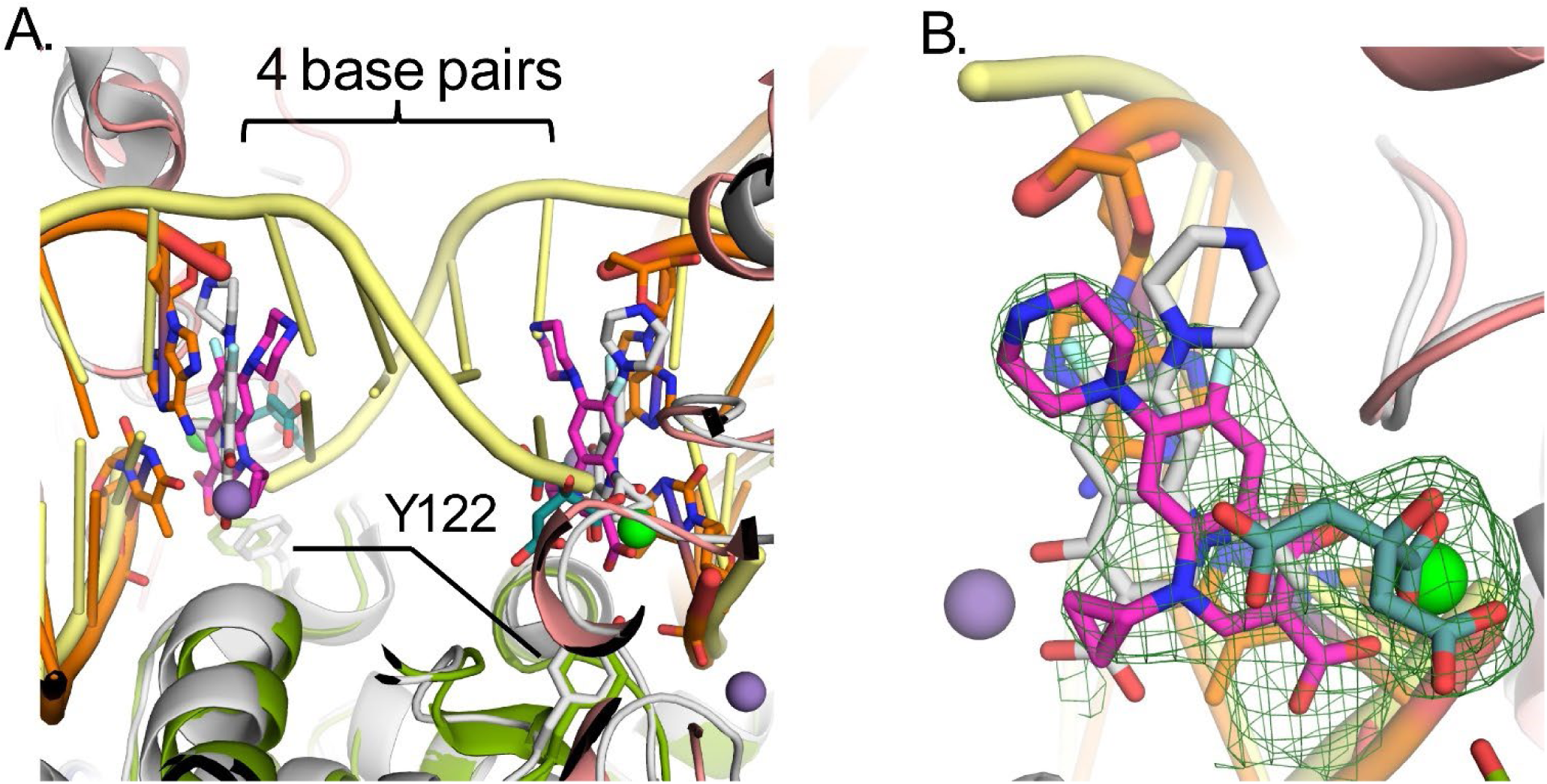
Comparisons of CPFX binding to DNA-bound gyrase structures in *P. aeruginosa* and *S. aureus*. (**A**) Active site superpositions with *S. aureus* protein (PDB ID 2XCT) [22] and CPFX-bound colored gray (manganese in purple) with DNA (yellow) to that of *P. aeruginosa* colored as in Figure 1. A citrate molecule (dark teal) co-chelates a manganese ion (green) with CPFX. For the *P. aeruginosa* structure, CPFX is binding stacking to the blunt ended DNA present at the active site. The DNA in *S. Aureus* binds as duplex DNA across the dimer interface with a nick on each strand of the DNA at the active site where CPFX intercalates the duplex DNA. A gap of four bases exists between the two active sites in each structure. The positions of the catalytic tyrosine residues of *P. aeruginosa* superimpose well with the inactive phenylalanine mutant of the *S. aureus*, supporting the protein is in a catalytically relevant conformation. (**B**) Face on view of CPFX binding in the two structures. Electron density from the Polder difference map for CPFX is shown contoured at 3.5 σ, displaying the difference in binding orientations between CPFX bound in the *P. aeruginosa* and *S. aureus* structures.

As seen for other gyrases such as in *E.* coli [30,38], structural alterations take place in *P. aeruginosa* gyrase upon binding DNA (Figure 3). Like in *E. coli,* the TOPRIM domain of GyrB undergoes a large change including C-terminal helix moving away from the active site allowing DNA to bind (Figure 3A). DNA binding also results in the insertion domain becoming mostly disordered (Figures 3B and 3C). Upon this movement, the Arg790 from the GyrB C-terminal helix becomes ordered and interacts with the DNA (Figure 3A, red asterisk). Residues on the loop between Ile452 and Asp461 become ordered as well, and shift to the position of the C-terminal helix in the apo structure with previously disordered residues Asn454 and Lys457 forming interactions with the DNA. The GyrA loop containing residues Ala175 to Met178 shifts to accommodate the DNA with the backbone amide of Gly177, forming a hydrogen bond to a DNA backbone phosphate. These changes further help to position the catalytic Tyr122 from the other molecule in the dimer, closer to the active site. The position of Tyr122 superimposes well with that of the inactive mutant Y123F of *S. aureus* [22], supporting it in a catalytically relevant position (Figure 4A). Residues Thr83 and Asp87, mutations that confer CPFX resistance, are found to be adjacent to the blunt end of the DNA and CPFX in this study (Figure 3A).

Although the DNA did not bind across the active site as expected, it did create a closed form of the enzyme that resembles other gyrases structures with regions of DNA and active site residues in positions consistent with catalytically relevant structures (Figure 2D and 4), generating a good starting model (Figure S2) for MD studies of the effects of mutations T83I and D87N on CPFX resistance.

### 3.2. T83I and D87N Mutations Do Not Drastically Change Structure of the Gyrase/DNA Complex or CPFX Binding

To gain mechanistic insight into *P. aeruginosa* CPFX resistance caused by T83I and D87N mutations in GyrA, we employed MD simulations to address the effects of these mutations on CPFX binding. We used six different systems (Materials and Methods, Table S1). The wild-type (WT) system was generated to serve as a reference; both GyrA molecules in the heterotetrameric system were mutated to generate either the T83I or the D87N mutant. Each of the three systems were subjected to MD in triplicate with and without CPFX bound at both active sites. To validate that these systems were dynamically stable, the Root Mean Squared Deviations (RMSD), a measure of the movement of atoms within a structure compared to those within a reference structure, were assessed. RMSDs of all backbone atoms stayed below 4 Å (Figure S3), except in the case of the CPFX-free T83I mutant (Figure S3B), for which the final values were slightly above 4 Å in one of three simulations. In most cases, GyrB has larger RMSDs than GyrA, mainly due to the contributions from large deviations culminating from the termini of two peptide fragments (residues 405-561 and 736-798) representing GyrB. Overall results indicate that the protein assemblies during simulations were stable on a global scale with or without bound CPFX during our simulation timescale (1000 ns). All CPFX-bound structures displayed slightly reduced RMSDs compared with their ligand-free counterparts (Figures S3).

CPFX, in its bound conformation, has its quinoline ring tightly stacked between the four DNA bases at the nick site (Figure S2). Accordingly, all RMSD values of the heavy atoms of CPFX are found to be close to 0.5 Å at the beginning of each production run and remain low throughout, suggesting that CPFX is tightly bound (Figure S4). Owing to the free rotation of the piperazine group around the connecting N-C bond to the quinolone core, the RMSDs can increase to values around 1Å for some systems during molecular dynamic simulation time frame (Figure S4). On the average, it displayed a deviation of about 0.75 Å in the center of mass position (Figure S4), even after this rotation of the piperazine group was considered. Still, this represents a rather low deviation, given the fact that even the smallest Root Mean Squared Fluctuations (RMSF) for individual backbone atoms of the amino acid residues of GyrA and GyrB surrounding the active site fall within this range (Figure 5). RMSF values represent the local fluctuations of each amino acid residue, and our MD runs do not show obvious difference between the RMSFs of the CPFX-bound and -unbound forms for the WT system or mutant systems (Figure 5). This stability of CPFX in the bound form may be attributed to the initial placement and orientation of CPFX that was based on the X-ray crystal structure of gyrase from *Mtb* with the complete DNA segment present (PDB ID 5BTC) [24]. The RMSF analysis show that WT and mutant systems behave in a very similar way, irrespective of the presence of CPFX (Figure 5). This suggests that these mutations have little effect on the local folding of CPFX-bound structures and, therefore, change in stability of CPFX-bound mutant complexes would likely not be a factor contributing to acquisition of antibiotic resistance.

**Figure 5.**
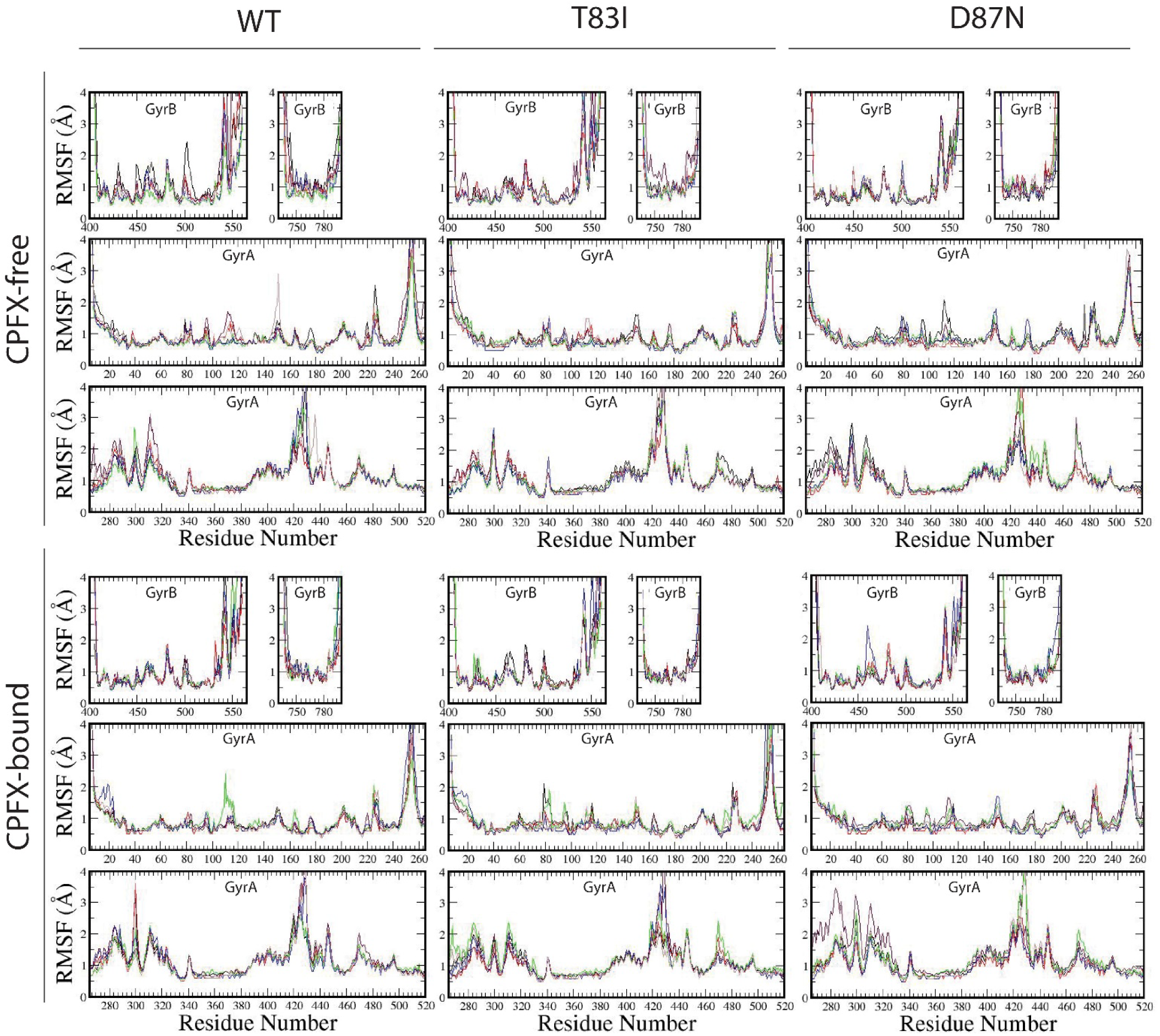
CPFX binding does not alter the local structure of the WT or mutant- systems as assessed by Root Mean Squared Fluctuations (RMSF) analysis. RMSF values of the WT, T83I, and D87N CPFX-free (top panels) and CPFX-bound (bottom panels) systems are shown. RMSF values were calculated by averaging the fluctuations of backbone atoms at the residue level. Values around 1 Å represent the stable, structurally important groups of residues while the larger fluctuations emanate from the loop areas. CPFX-binding does not alter the local structures of any system.

Dynamic Cross-Correlation (DCC) analyses of alpha carbons, employed to search for correlated motion between residues revealing functional networks, clearly indicate that, in the WT system, the segment of the GyrA directly in contact with the segment of GyrB or DNA has highly positive correlations, while the non-interacting portions (residues 350-513 of GyrA) distinctly display their own dynamics. This is true for each of the GyrB/GyrA complexes (I and II) (Figure 6). Further analysis of the cross-correlation behavior between complexes I and II reveals similar but much weaker correlation (Figure 6, top and middle right panels). The coiled-coil region (residues 348-354) of GyrA shows the least correlation with the adjacent complex in each system except for the gate residues at the base of the structure (Figure 2C; ∼residues 390-470), which strongly correlates with the adjacent complex’s gate residues due to the surface contacts. No appreciable differences between the CPFX-free and -bound forms of the WT system points to having similar dynamical behavior irrespective of the presence of the drug (Figure 6, bottom panels). The same tendencies in the pattern of interactions were observed for both mutant systems (Figure 7). These data suggest that dynamics of interactions between GyrA and GyrB in mutant complexes do not differ significantly from the WT complexes. Moreover, collinearity of the protein dynamics of these complexes with or without the bound ligand CPFX, either in mutant or WT systems, argues that such dynamics does not underlie antibiotic resistance of mutant complexes.

**Figure 6.**
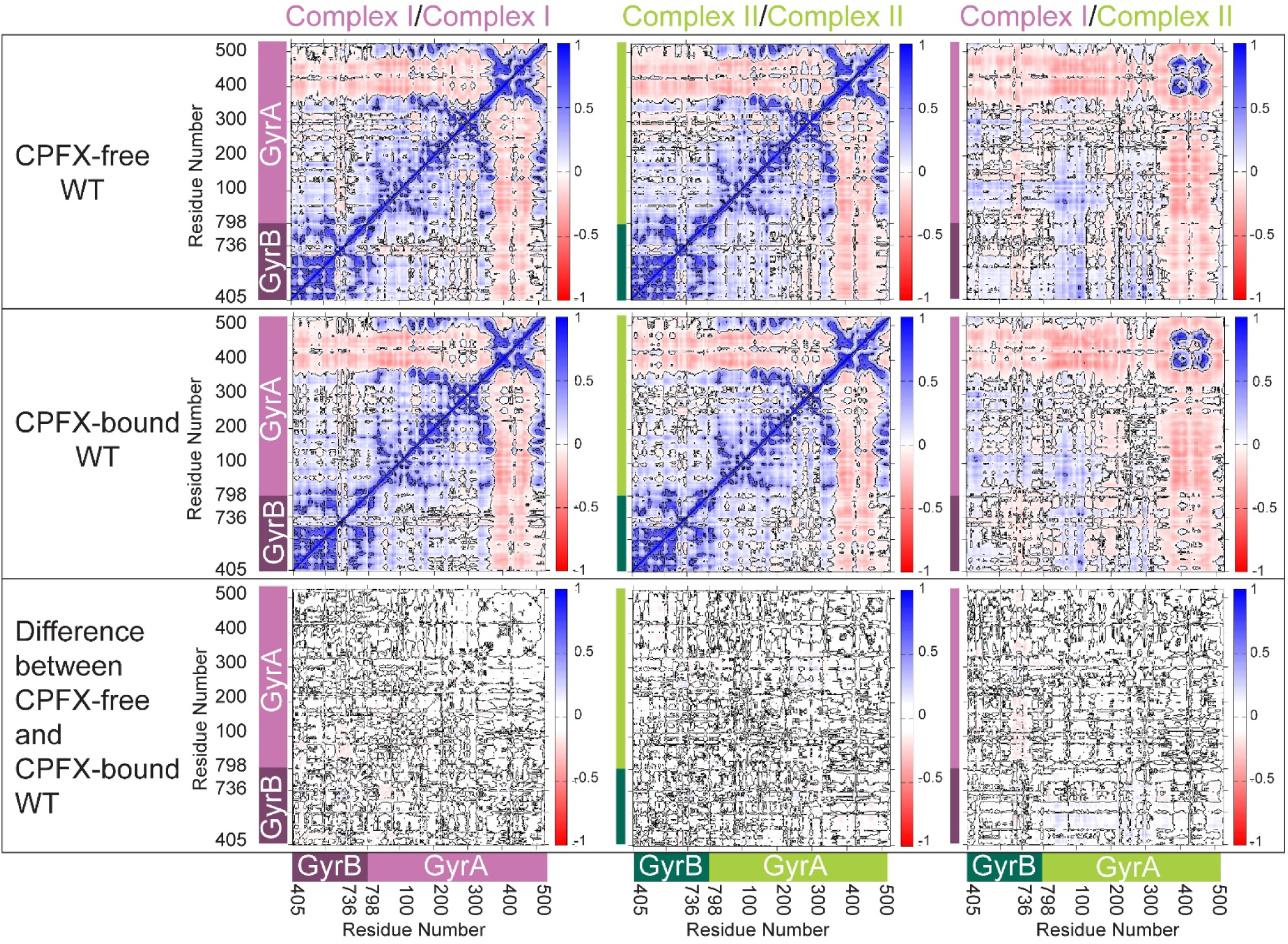
CPFX binding does not alter dynamic correlations in the WT system. Dynamic Cross-Correlation Matrices (DCCMs) display that molecules or domains move similarly against each other during MD simulations. DCCMs were calculated using the positions of alpha carbon atoms and averaged over a microsecond for the three runs executed for each system. The top row shows values of DCC for the CPFX-free system. The middle row shows values of the CPFX-bound system. The bottom row shows the difference DCCMs between the CPFX-free and CPFX-bound systems. The left and middle columns show the DCMMs of complexes I and II, respectively. The right column shows the values of cross-correlations between complex I and II within the tetramer system. Correlated motion is depicted by blue patches, no correlation by white, and the anti-correlated motion by red patches.

**Figure 7.**
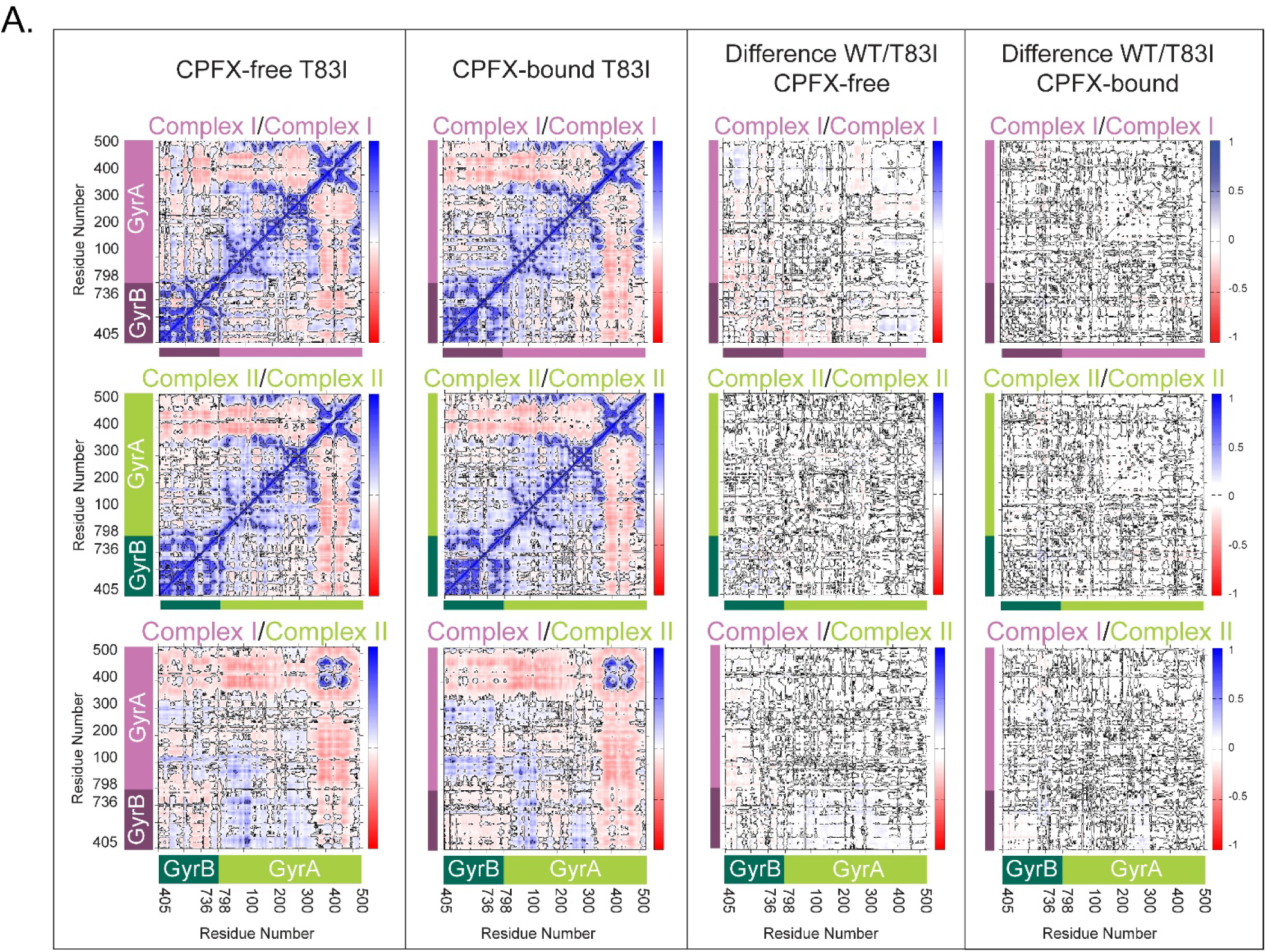
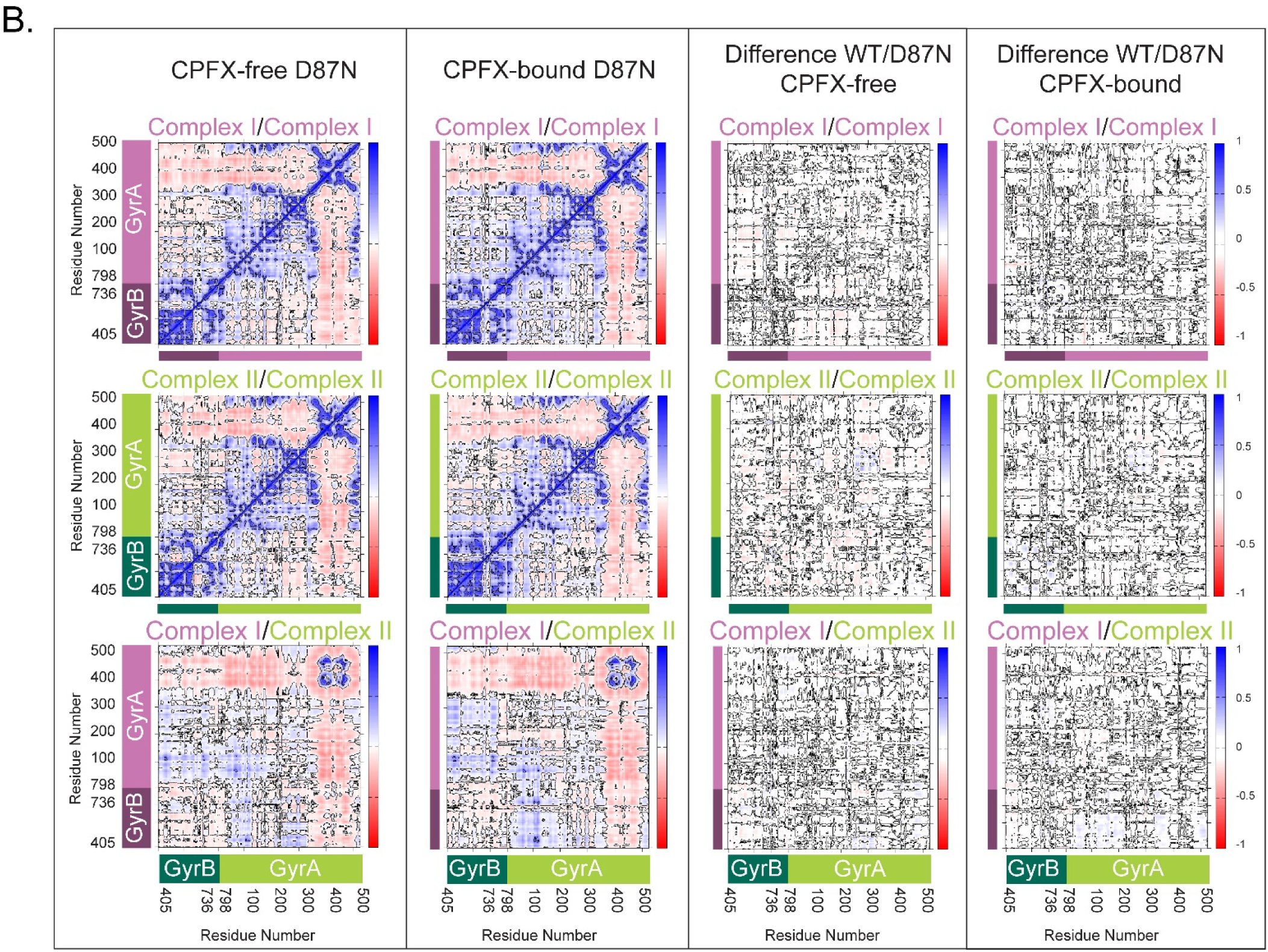
Dynamic correlations in mutant complexes do not differ significantly from the WT complexes. DCCMs were calculated using the positions of alpha carbons averaged over a microsecond for three runs for each system. The left column shows values of DCCM for the CPFX-free state. The second column from the left shows values for the CPFX-bound mutant form. The second column from the right shows the DCCM differences between the CPFX-free mutant and WT, while the rightmost column is for the CPFX-bound mutant and WT difference. The top and middle rows show the dynamic auto cross-correlation matrices of complexes I and II, respectively. The bottom row shows the values of cross correlations of complexes I and II. The correlated motion is depicted by blue patches, no correlations by white, and the anti-correlated motion by red patches. Whether CPFX-bound or not, DCCM plots show strikingly similar behavior in their cross correlations with only minor differences in the difference matrix plots. (**A**) T83I, and (**B**) D87N.

Whether CPFX-bound or not, DCCM plots show similar cross correlations. GyrA (complex I in salmon pink; complex II in light green) and GyrB (complex I in maroon; complex II in green) are marked as side bars.

### 3.3. DNA Plays a Significant Role in Maintaining the CPFX-Bound Conformation

Molecular Mechanics with Generalized Born and Surface Area (MMGBSA) solvation energy calculations were used to quantify the residue (or nucleotide) contribution for CPFX interaction energy. Only the attractive interaction energies (negative values) are displayed (Figure 8A-C). Only residue Thr83 of WT GyrA exhibits appreciable, attractive binding interaction to CPFX with solvation energies of approximately -3 kcal/mol (Figure 8A, Thr83, white circles). In the T83I system, the interaction strength decreases by about 2 kcal/mol (Figure 8A, compare Thr83 WT (white circles) and T83I (red circles)). The interaction energy with Asp87 was undetectable and not shown. Surprisingly, a decrease in the interaction energy of Thr83 with CPFX is seen to a lesser extent even in the D87N system, indicating subtle local changes due to this mutation (Figure 8A, Thr83, blue circles). In addition, we report here a small but detectable interaction with GyrA Arg121. In fact, this residue belongs to GyrA of the heterodimer complex that is not primarily involved in CPFX interactions (Figure 8A, Arg121).

**Figure 8.**
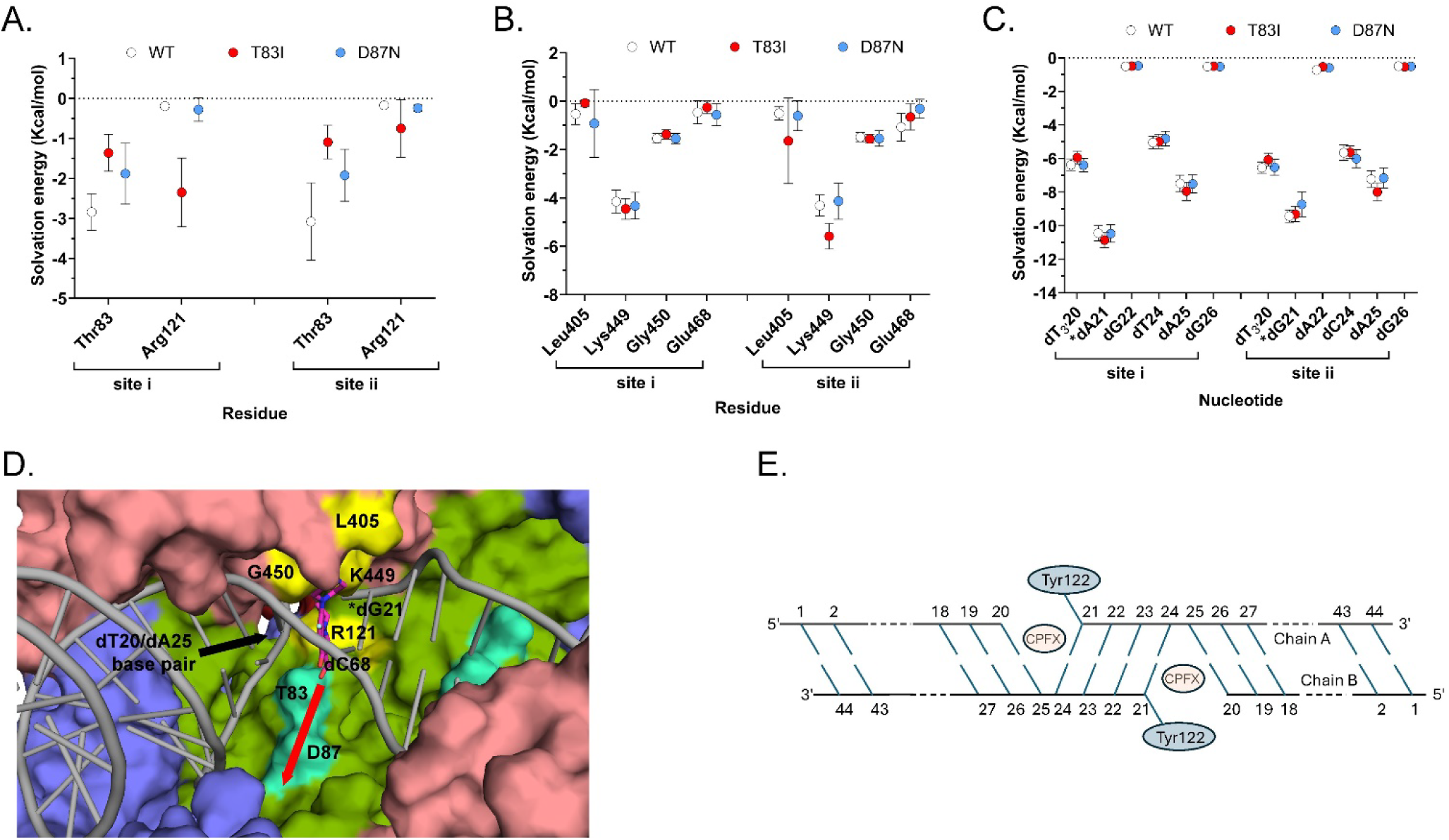
DNA plays the significant role in maintaining the CPFX-bound conformation. Graphs (**A**), (**B**), and (**C**) represent the energy of CPFX-residue interaction calculated for various simulated systems. (**A**) CPFX interaction energy with residues Thr83 and Arg121 of GyrA. (**B**) CPFX interaction energy with residues Leu405, Lys449, Gly450, Glu468 of GyrB. (**C**) CPFX interaction energies with nucleotides dT24, dA25, dG26, dT3′20, *dA21, and dG22, dT3′20, *dG21, dA22, dC24, dA25, and dG26 at the CPFX binding site where * indicates the nucleotide covalently bound to Tyr residues. site i and site ii are the two sites involved in CPFX binding as shown in Figure S2. Symbols represent mean +/- SE (n= 3). Image (**D**) represents the surface rendering of the water-filled cavity (referred to as the cavity-site) in GyrB/GyrA-GyrB/GyrA heterotetramer above residues Thr83 and Asp87. This cavity, detected in all simulations, may represent the path for CPFX to access the active site and intercalate between two pairs of nucleotides. CPFX was moved along the direction shown by the red arrow using MD with Umbrella Sampling to estimate the influence of residues Thr83 and Asp87 and their mutants have on the binding process. Displayed is one of the active sites colored as in Figure 1A but with residues Thr83 and Asp87 in green-cyan, and Leu405, Lys449, G450 and Arg121 (from the other GyrB/GyrA-GyrB/GyrA dimer) in yellow. Glu468 is hidden behind other residues in this figure and is not visible. (**E**) Molecular schematic of the DNA nucleotides included in 8C and D.

Several residues of GyrB contribute to CPFX interactions, with Lys449 standing out as particularly prominent across all systems due to its proximity to CPFX (Figures 3A and 8B). Other interacting residues, including Leu405, Gly450, and Glu468, show reduced interaction strengths of one or more kcal/mol (Figure 8B) compared to Lys449. Based on the MMGBSA energy values of DNA nucleotide interactions with CPFX (Figure 8C), we concluded that the DNA plays the most significant role in trapping CPFX between the nucleotide bases. As shown in Figure 8D and E, the most significant interaction energy contribution comes specifically from the nucleotide bases directly engaging with CPFX, while neighboring nucleotides exhibit only residual interactions. Interestingly, CPFX in the crystal structure (Figure 4) is found binding against the equivalent base to dA25 in the simulation (Figure 8D and E), which shows the greatest contribution to binding (Figure 8C). This suggests its presence is enough to energetically bind CPFX, but base dC24 of the four-nucleotide gap helps to properly position CFPX as seen in other crystal structures and utilized for this simulation [22,24].

### 3.4. Reverse MD Suggests Importance of a Solvent Cavity Adjacent to Thr83 and Asn87 for CPFX Access

Since we did not find compelling evidence that the mutations T83I or D87N have a significant effect on gyrase structure or CPFX-binding dynamics, we focused on further investigating the dynamic process of CPFX binding rather than on the bound conformation itself. Notably, all structural analyses reveal a water-filled cavity adjacent to the bound-CPFX that sits directly adjacent to both Thr83 and Asp87 (Figure 8D). The cavity is formed around the residue segment Asp82-Asp87 and residues Val90, Arg91, Gln94, Phe96, Ser97, Asn116, Ala117, Met120, Asn268, Asn270, Ala272, and Arg273, along with the backbones of the residue segment Ser111-Asp115, and here after referred to as the “cavity site”. While CPFX does not directly interact with Asp87 (Figures 3A and 8D) in the bound form, we believe that the cavity site might offer insights into how Asp87 influences CPFX’s positioning between nucleotides for its function.

We hypothesize that in its path to the inhibition binding site, CPFX first diffuses to the cavity site from its solvated state, before it finds its destination between the nucleotide bases. There is an additional small cavity that could act as another channel to enter the inhibition site located right above the four nucleotides that sandwich CPFX. However, the net negatively charged CPFX would have to travel near the DNA backbone, if it were to use this as the entry point. The small size of this cavity and the negative charge likely work against utilizing this second site, and therefore, we did not further investigate CPFX binding via this cavity.

To explore CPFX access to the cavity site, we performed a reverse experiment, using MD and the Umbrella Sampling technique to guide CPFX into the mentioned cavity from its binding site, path that is illustrated in Figure 8D. Specifically, we steered CPFX away from its bound position and constrained it within a window of 0.4 Å thickness, applying forces to its atoms. Each window was sampled for 1.2 ns, and 500 samples, collected from the final nanosecond of each window, were subjected to MMGBSA energy analysis to determine the interaction energies of CPFX with Thr83 or Ile83 and Asp87 or Asn87.

Figure 9 illustrates the changes in distance between the oxygen atoms in CPFX and the gamma-oxygen of Thr83 and Asp87 over time as shown by a representative Umbrella Sampling trajectory.

**Figure 9.**
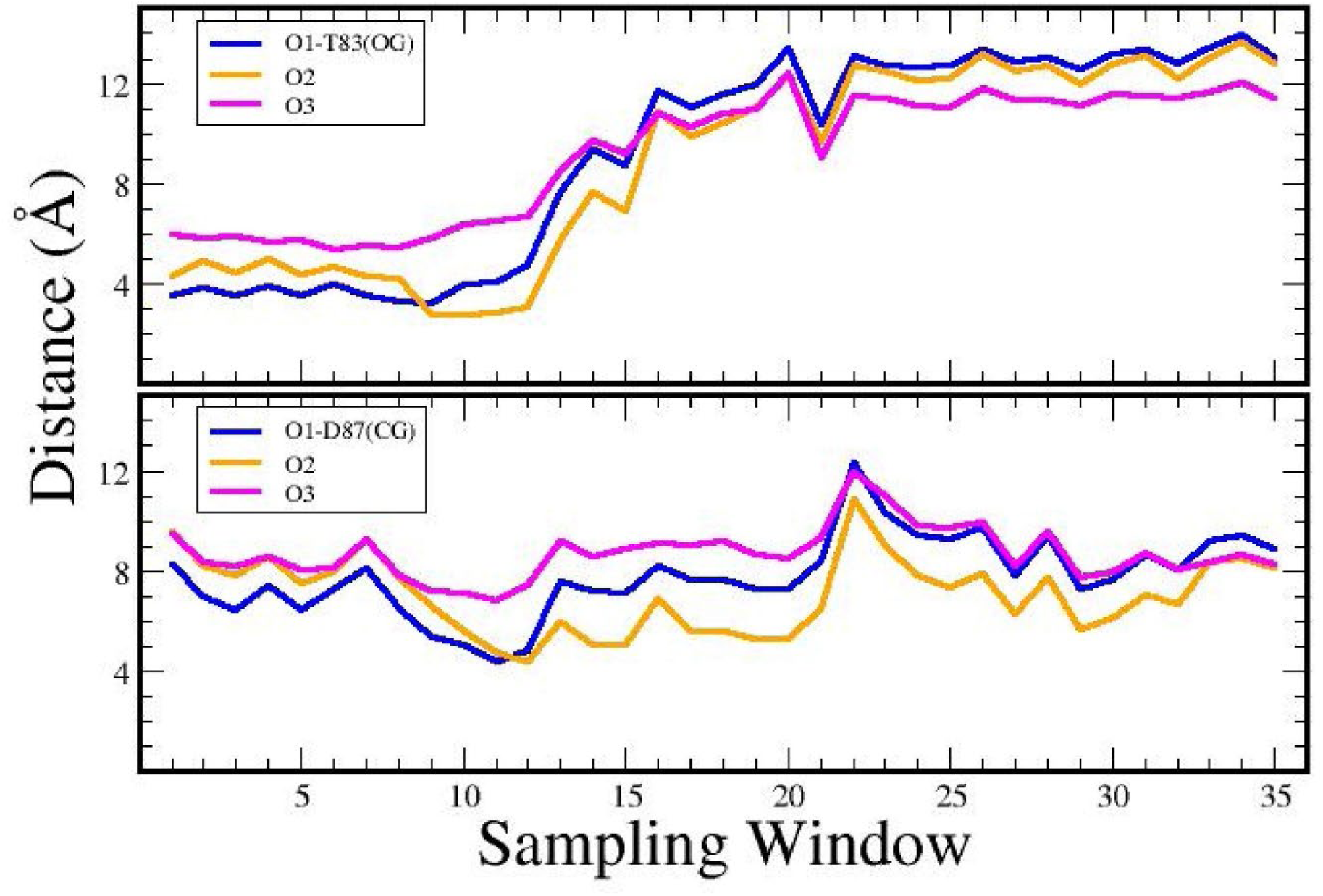
Distance plot of CPFX atoms to Thr83 and Asp87 when moved out of binding site. Distances were calculated for the movement of CPFX from the bound position to the cavity described in Figure 8A. The three oxygens of CPFX to residues Thr83 (gamma oxygen “OG” was used in measuring distances), shown in the top graph, and Asp87 (gamma carbon “CG” was used here), shown in the bottom graph, during MD with Umbrella Sampling.

Initially, Thr83 was in close contact with one or more oxygen atoms (sampling windows 1-12). Later, Thr83 and CPFX separated over the course of the simulation. In contrast, the distal gamma-carbon of Asp87 moved closer to one or more CPFX oxygen atoms within the same trajectory, shifting to CG (Gamma Carbon) distances around 5 Å away from its original bound state for a set of sampling windows (10 to 20).

MMGBSA energy analysis shows that, in the WT system (Figure 10A), Thr83 presents a distinct attractive center for CPFX within a range of 0–5 Å from its inhibitory binding state, with an interaction strength between 2–5 kcal/mol. Conversely, Asp87 exhibits a clear attraction to CPFX at a distance from its inhibitory binding state, with interaction magnitudes ranging from approximately 1–5 kcal/mol. The combined effects of these interactions may facilitate CPFX’s entry into the cavity site before it becomes locked between the two sets of nucleotide bases in the DNA.

**Figure 10.**
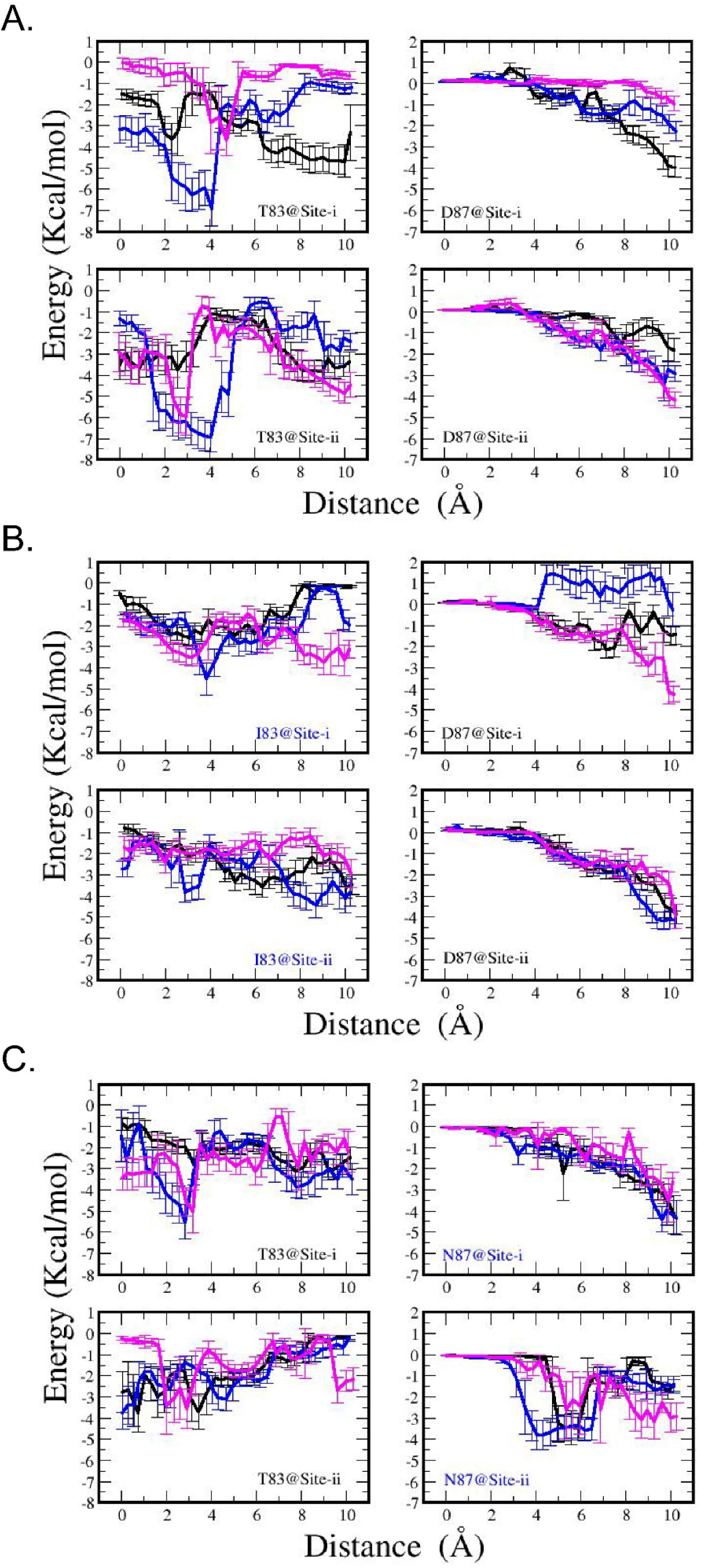
Thr83 provides a favorable environment for CPFX at or near its binding site. Changes in interaction energy between CPFX and Thr83 or Asp87 (and mutant residues) as CPFX is moved along the trajectory from the inhibitory binding site to the adjacent cavity-site entrance during Umbrella Sampling MD calculated for; (**A**) WT, (**B**) T83I, and (**C**) D87N systems. Each color corresponds to one umbrella sampling trajectory. Since the entrance to the cavity site is non-uniform, similar nonuniform energetics can be found in most systems. However, a clear signature of a favorable interaction of -3 to -7 kal/mol about 2 to 4 Å away for the CPFX inhibitory binding position is observed for Thr83. Once mutated to Ile83, the interaction energy drops below -4 kcal/mol. For Asp87, a stronger interaction is observed for CPFX distal to the inhibitory binding site. This behavior appears altered in both mutant systems.

In the T83I mutant system (Figure 10B), there is a suppression of the attractive well observed for Thr83 in the WT system at 4 Å distance, accompanied by modifications of the Asp87-CPFX interactions at as well. On the other hand, in the D87N system (Figure 10C), the Asn87-CPFX interactions are somewhat stronger across a broader distance range. Interestingly, Thr83-CPFX interactions within this complex are also altered (compared to Figure 10A, left panels), showing a weaker attractive potential than in the WT complex.

Since the cavity site may provide access to the inhibitory bound state of CPFX, our findings argue that the T83I mutation may have a direct effect on the process of CPFX entering the inhibitory binding site, with its greatest contribution when the drug has moved 2 to 4 Å from its bound state. This favorable interaction diminishes when Thr83 is mutated to isoleucine (Figure 10B). For Asp87, the interaction energy is the greatest when CPFX has moved from the inhibitory position to the cavity site. As CPFX progressed further into the cavity and approached Asp87, the interaction energy in the WT system increased (Figure 10A).

## 4. Discussion

*P. aeruginosa* represents a clinically relevant target for new antibiotics due to its prevalence, severity of infection outcomes, and intrinsic, acquired, and adaptive resistance to antibiotics [51,52]. Two mutations in the DNA gyrase that have been shown to render *P. aeruginosa* resistant to fluoroquinolones such as CPFX are T83I and D87N [10,12,15]. However, how these mutations contribute to *P. aeruginosa* resistance is not well understood. To address this clinically relevant question, we structurally characterized the catalytic core of the WT gyrase relevant to CPFX resistance and performed molecular dynamics (MD) to evaluate the effect that these mutations might have on CPFX binding.

The apo crystal structure showed good agreement with the previously determined structure of the *E. coli* gyrase except for a shift in the position of the insertion domain (Figure 2B). DNA binding results in conformational changes surrounding the active site to accommodate the DNA and catalytically relevant positions of Tyr122 residues for DNA cleavage. Surprisingly, in our study, the DNA binds with two blunt ends in the active site. This structure may mimic that of the complex post-melting of the four DNA base pairs between the two nicks in the G-segment, leaving a 4-base pair gap through which the T-strand would pass to supercoil the DNA. The positions of the DNA and catalytic Tyr122 are in good agreement with the configuration of the equivalent residues reported for other gyrase structures [22,24,30]. CPFX is also found binding at the same position as in the *S. aureus* and *Mtb* structures although the relative orientation differs, potentially due to the lack of the stacking base pair on the gap side (Figure 4A) [22,24].

MD simulations have previously been employed to investigate delafloxacin (also a fluoroquinolone) resistance in *S. aureus* gyrase [28], revealing that the mutations studied (S84L/E88K, S84L/S85P, and single mutation S84L) do not induce significant structural perturbations upon drug binding. Notably, the wild-type gyrase exhibited the most negative binding free energy, suggesting it offers the most favorable environment for delafloxacin interaction. The study further shows that the mutations increased the plasticity within the CPFX-binding pocket and elevated the atomic mobility of the mutated proteins, as demonstrated by principal component analysis. Interestingly, residues Ser84, Ser85, and Glu88, located within the fluoroquinolone-binding pocket (residues Thr83 and Asp87 in *P. aeruginosa* correspond to Ser84 and Glu88 in *S. aureus*) were found to neither directly interact with delafloxacin nor significantly contribute to its binding energy [28]. However, the absence of DNA in the MD simulations performed in the quoted study may give an inaccurate description of the inhibitory site, since CPFX is found sandwiched between DNA bases in the gyrase crystal structure with DNA bound [22,24].

Using the GyrA crystal structure of *Colwellia psychrerythraea*, Sada et al. [15] constructed a homology model of the *P. aeruginosa* GyrA along with the GyrA T83I and the GyrA D87N single mutants, and the double mutant GyrA T83I/D87N, and performed a docking study with CPFX. The affinities calculated directly from the docking poses using the GyrA systems resulted in very similar binding energies for the WT and mutant systems (-6.98±0.42, -6.2±0.19,-6.9±0.46, and -6.5±0.52 kcal/mol for the WT, T83I, D87N, and T83I/D87N systems, respectively), with only a slight favor for the WT. Our study, based on an X-ray crystal structure of *P. aeruginosa* GyrB/GyrA-GyrB/GyrA catalytic core with bound DNA and CPFX, suggests that GyrB residues such as Lys449 (Figure 3A) can also influence CPFX binding and thus should be included when evaluating quinolones binding. When GyrB was included in the analysis, the architecture of the quinolone binding cavity was redefined: the floor was assigned as residues 87–99 of GyrA and the roof as residues 447–449 of GyrB. Moreover, the residues 67–106 of GyrA —collectively known as the Quinolone Resistance-Determining Region (QRDR)— were confirmed as a critical mutational hotspot associated with resistance development [53].

In the present study, we have shown that the mutations T83I and D87N, located in the QRDR, do not appear to cause notable structural or dynamical perturbations within the *P. aeruginosa* gyrase complex. Global stability, as measured by Root Mean Squared Deviations (RMSD), and local conformational flexibility, assessed through Root Mean Squared Fluctuations (RMSF), remained largely unaffected by either drug binding or mutation (Figure 5, S3 and S4). Cross-correlation analyses highlighted a pronounced association between GyrA and GyrB subunits, while inter-monomer correlations were markedly lower than the autocorrelations observed within each individual monomer. A notable exception was found in the exit gate residues of GyrA, which exhibited enhanced correlated motion across the two monomers, indicating a region of increased cooperative dynamics (Figures 6 and 7).

MD simulations rendered insight into a solvent-filled cavity, preserved in all simulated systems, that may provide access to the inhibition site (Figure 8D), offering an opportunity to investigate the reverse pathway —namely, CPFX entry into its inhibition site via this cavity site. By applying a sampling technique commonly used in MD (Umbrella Sampling) and guiding CPFX along a pre-defined path back towards the cavity site, we were able to generate data informative for studying the reverse process of CPFX binding.

In the bound state, residue Thr83 appears to be in proximity (<5 Å) with the drug, while Asp87 remains spatially distant (sampling window 0, Figure 10A). Energy profile analyses of the WT system (Figure 10A) suggest that Thr83 contributes favorably to the interaction energy in the initial stages of CPFX extraction from the binding site, with its greatest contribution when the drug has moved 2 to 4 Å from its bound state. This favorable interaction diminishes when Thr83 is mutated to isoleucine (Figure 10B). For Asp87, the interaction energy is the greatest when CPFX has moved from the inhibitory position to the cavity site. As CPFX progressed further into the cavity and approached Asp87, the interaction energy in the WT system increased.

## 5. Conclusion

Our results suggests that while the mutant residues do not render an increase in CPFX binding energy (decreased affinity) in the bound state, they may influence the actual access of CPFX to the inhibitory site by reducing the attractive forces associated with the process of binding. These findings underscore the complexity of molecular interactions, in which even subtle, non-destabilizing mutations may alter the trajectory of CPFX entrance to the active site. Understanding the importance of the path by which CPFX enters the active site could not have been elucidated solely by analysis of static (crystal or cryo-EM), but became possible in combination with modelling of the interaction dynamics. We believe that combined structural analysis and MD simulations that consider antibiotic access may be a useful consideration for screening compound libraries to identify new therapeutic agents to target fluoroquinolone resistant bacteria.

## Acknowledgements

We thank Dr. Yukitomo Arao for critical reading of the manuscript .

## Supplementary Materials

Figure S1: CPFX-induced cleavage reactions of gyrB397-GGS-gyrA525; Figure S2: Space filling diagram of the modeled WT GyrB/GyrA tetramer system bound to CPFX and DNA; Figure S3: Root mean squared deviations (RMSD) show global stability of simulated systems; Figure S4: CPFX heavy atoms fluctuations in the gyrase active site as measured by RMSDs; Table S1: The composition of various gyrase systems (with and without CPFX) studied using molecular dynamic simulations (each run is 1 μsec).

## Author Contributions

L.P.: Methodology, Investigation, Data curation, Formal analysis, Writing—original draft; L.G-V.: Conceptualization, Methodology, Investigation, Writing—original draft; A.M.K.: Investigation, Writing—review and editing; N.D.: Conceptualization, Supervision, Writing—review and editing; L.C.P: Methodology, Investigation, Data curation, Formal analysis, Writing—original draft; P.W.D.: Resources, Project administration, Writing—review and editing. All authors have read and agreed to the published version of the manuscript.

## Institutional Review Board Statement

N/A.

## Informed Consent Statement

N/A.

## Data Availability Statement

The original contributions presented in this study are included in the article/Supplementary Materials.

