## Supplementary material for "Structural and Computational Analysis of *Pseudomonas aeruginosa* DNA Gyrase Reveals Molecular Characteristics That May Contribute to Ciprofloxacin Resistance": Figure S1

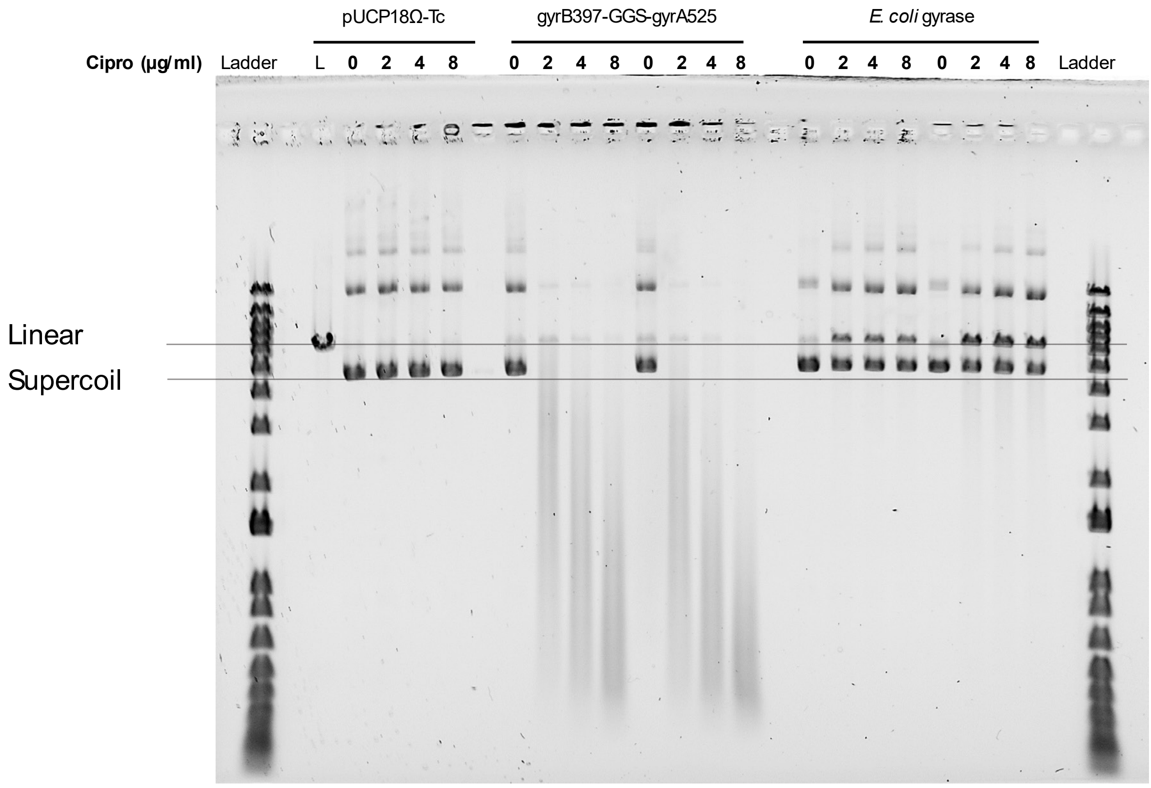


**Figure S1.** CPFX-induced cleavage reactions of gyrB397-GGS-gyrA525. Samples of a pUCP18 plasmid preparation (0.2 µM) were incubated with gyrB397-GGS-gyrA525 (1 and 2 µM) and *E. coli* gyrase (Sigma; 1 and 2 µl of a 2 units/µl solution) in the presence of different CPFX concentrations (0, 2, 4, and 8 ug/ml) for 1 hour at 37°C. After addition of SDS and proteinase K, and incubation for 30 minutes at 37°C, samples were analyzed on a 0.8% agarose gel. Plasmid integrity decreases with greater concentrations of CPFX in the presence of gyrB397-GGS-gyrA525, which indicates that the antibiotic inhibits the ligase activity of the construct after it cleaves the dsDNA plasmid. L stands for linearized.

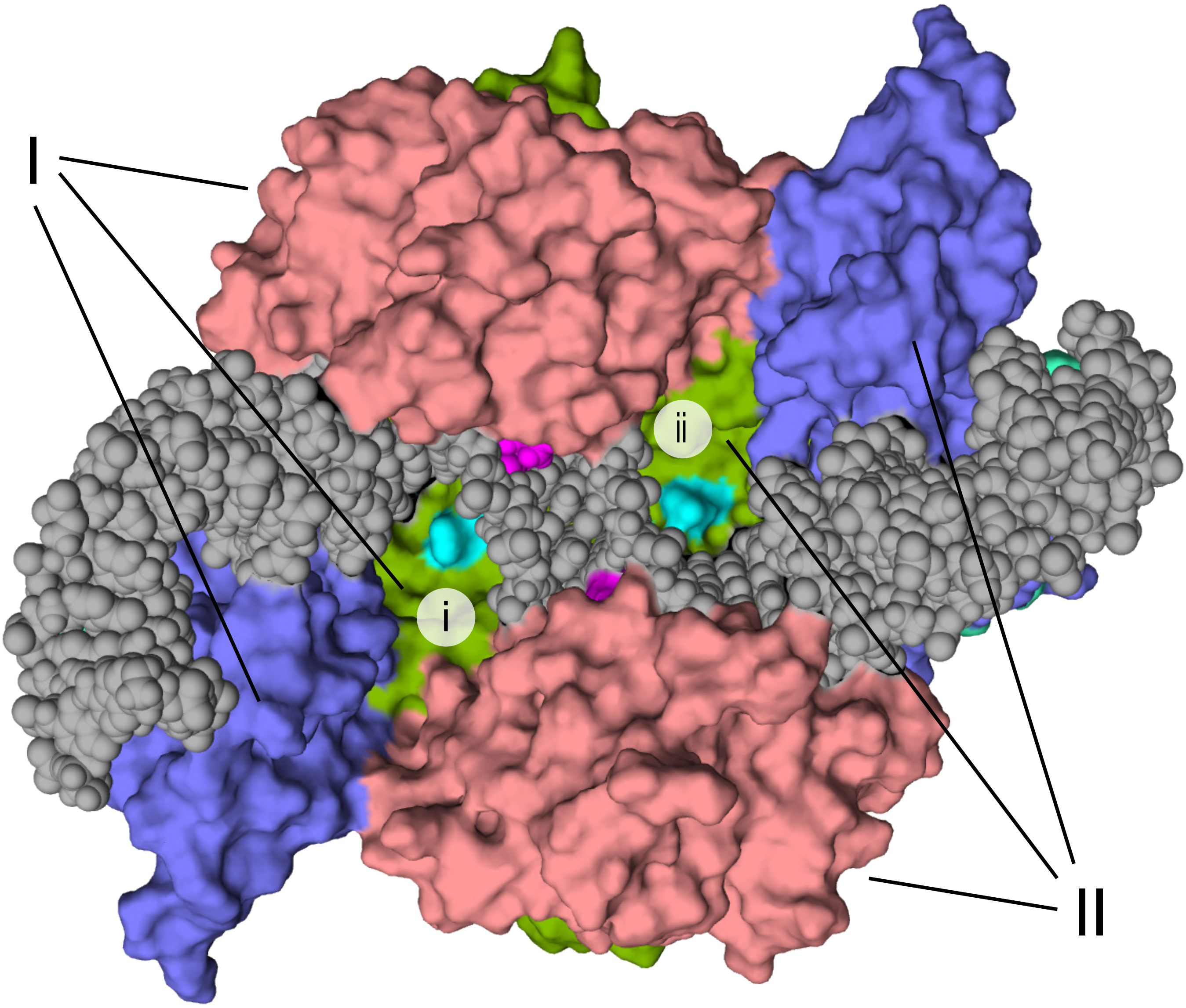


**Figure S2.** Space filling diagram of the modeled WT GyrB/GyrA tetramer system bound to CPFX and DNA. GyrB/GyrA complex I and II with DNA (atoms in gray spheres) and CPFX (atoms in magenta) form the model system. Gyrase is colored as in Figure 1A.

**Figure S3.** Root mean squared deviations (RMSD) show global stability of simulated systems. RMSD of the A)  WT, B) T83I, and C) D87N systems. RMSD values were calculated by superimposing the backbone heavy atoms of all residues with those at the initial conformation. Left panels show the overall RMSD of the entire system (Complex I and II, black) and each individual complex (denoted by I (red) and II (green)). On the right  side, RMSD are displayed for individual molecules (Gyrase A and B) of each complex (I and II). Triplicates correspond to the three simulations carried out on each system. The plateau in the values indicates each system has equilibrated rather well and is stable. CPFX-binding does not seem to alter the stability of any system at the global scale. Frequencies (number of samples with a certain RMSD value) were calculated to further analyze for the presence of  potential multiple conformations within a trajectory.

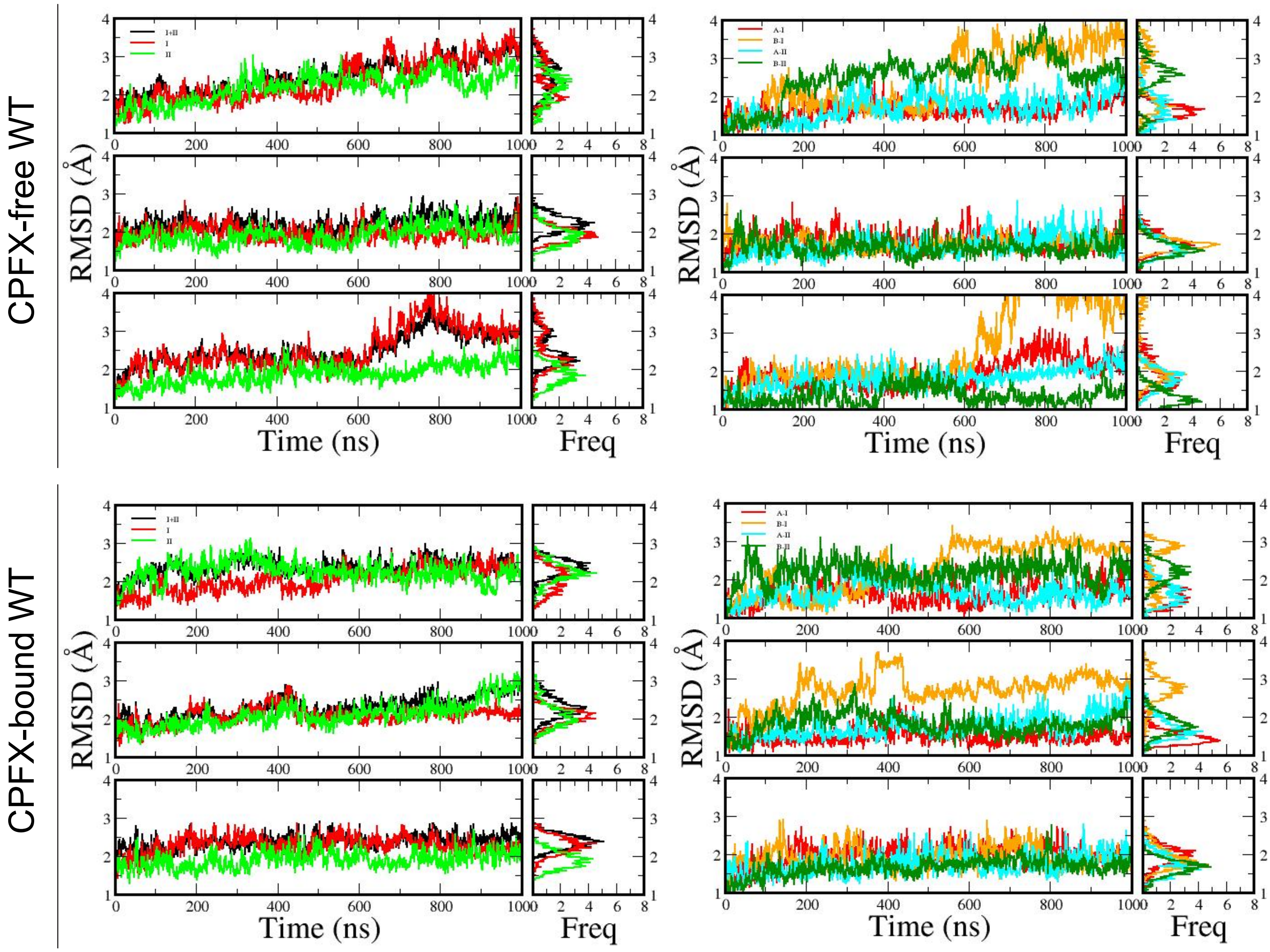


**Figure S3A.** Root Mean Squared Deviations (RMSD) of the WT system versus time (1000 ns).

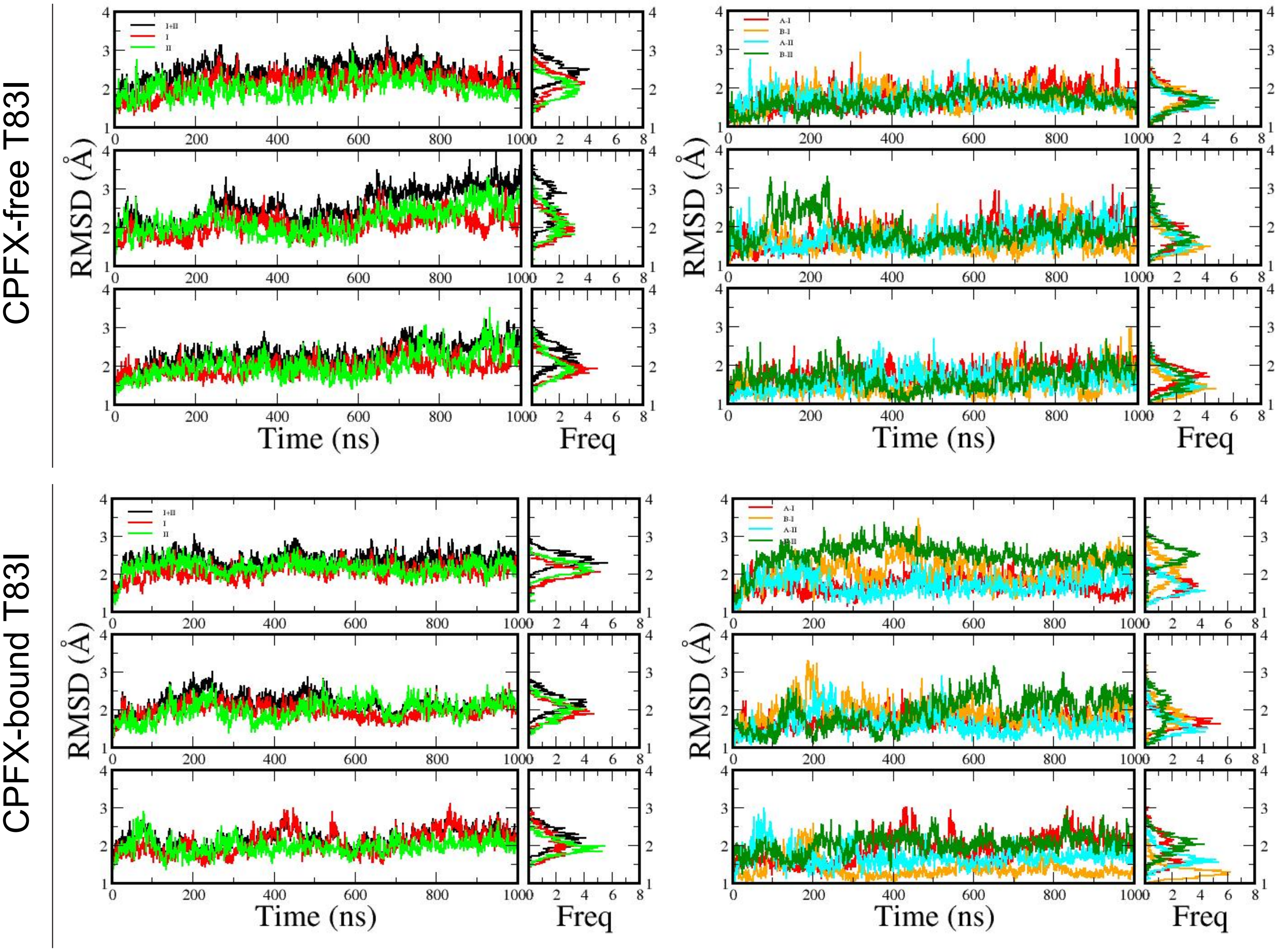


**Figure S3B.** Root Mean Squared Deviations (RMSD) of the T83I mutant system versus time (1000 ns).

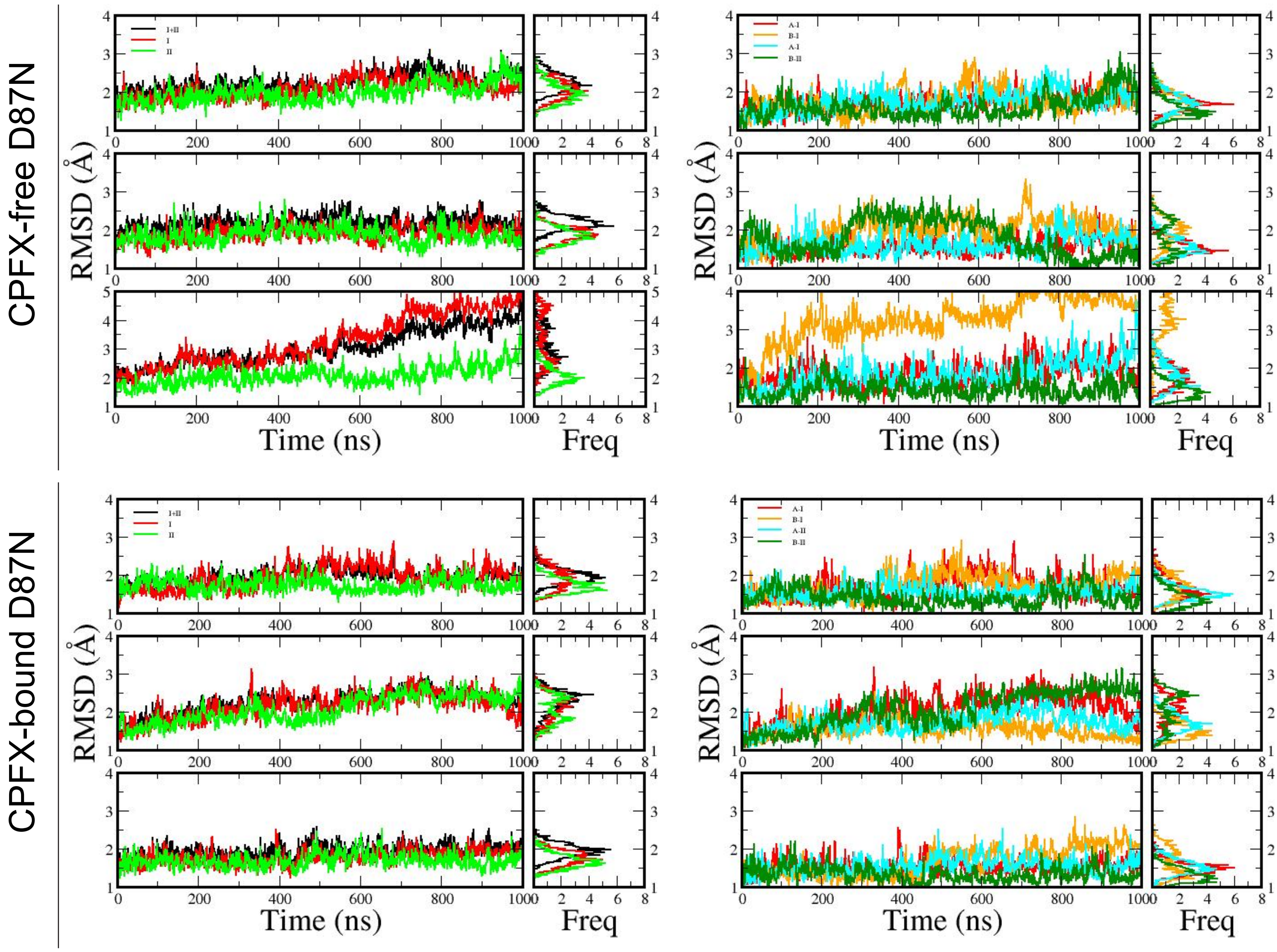


**Figure S3C.** Root Mean Squared Deviations (RMSD) of the D87N mutant system versus time (1000 ns).

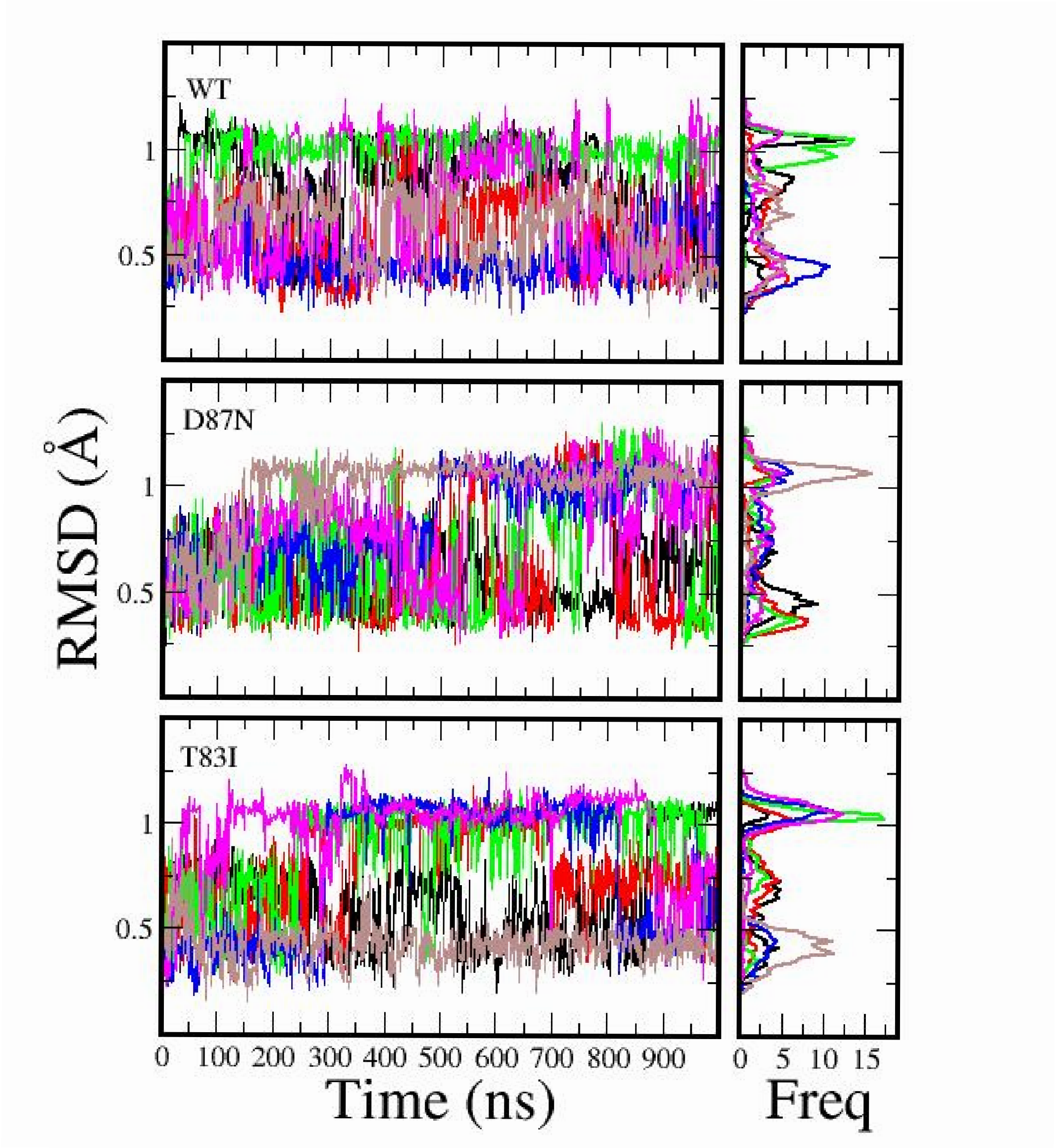


**Figure S4.** CPFX heavy atoms fluctuations in the gyrase active site as measured by RMSDs. Starting position was taken as the reference. Very little movement is observed for CPFX, suggesting it is tightly bound. A little over 0.5 Å variation is solely due to the free rotation of the piperazine group around the connecting N-C bond to the quinolone core. Each panel has 6 graphs shown in various colors corresponding to three MD runs and the two CPFX molecules in each run.

**Table S1**. The composition of various gyrase systems (with and without CPFX) studied using molecular dynamic simulations (each run is 1 $\mu$s).

| System | # of MD runs | Atoms | $Cl^-$ | $Na^+$ | Water molecules |
|---|---|---|---|---|---|
| WT | 3 | 279466 | 231 | 357 | 84266 |
| WT/CPFX | 3 | 279472 | 231 | 359 | 84200 |
| T83I | 3 | 279476 | 231 | 357 | 84266 |
| T83I/CPFX | 3 | 279482 | 231 | 359 | 84200 |
| D87N | 3 | 279468 | 231 | 359 | 84266 |
| D87N/CPFX | 3 | 279474 | 231 | 357 | 84200 |
